# BTSP, not STDP, Drives Shifts in Hippocampal Representations During Familiarization

**DOI:** 10.1101/2023.10.17.562791

**Authors:** A.D. Madar, C. Dong, M.E.J. Sheffield

## Abstract

Synaptic plasticity is widely thought to support memory storage in the brain, but the rules determining impactful synaptic changes in-vivo are not known. We considered the trial-by-trial shifting dynamics of hippocampal place fields (PFs) as an indicator of ongoing plasticity during memory formation. By implementing different plasticity rules in computational models of spiking place cells and comparing to experimentally measured PFs from mice navigating familiar and novel environments, we found that Behavioral-Timescale-Synaptic-Plasticity (BTSP), rather than Hebbian Spike-Timing-Dependent-Plasticity, is the principal mechanism governing PF shifting dynamics. BTSP-triggering events are rare, but more frequent during novel experiences. During exploration, their probability is dynamic: it decays after PF onset, but continually drives a population-level representational drift. Finally, our results show that BTSP occurs in CA3 but is less frequent and phenomenologically different than in CA1. Overall, our study provides a new framework to understand how synaptic plasticity shapes neuronal representations during learning.

## INTRODUCTION

Since Donald Hebb’s influential postulate (Brown & Milner, 2003; Hebb, 1949), learning and the encoding of memories are assumed to be mainly supported by activity-dependent synaptic plasticity (Dringenberg, 2020; Moldakarimov & Sejnowski, 2017). The dependencies of long-term plasticity (LTP) on neuronal activity have been studied for decades, but mostly in-vitro (Chistiakova et al., 2015; Feldman, 2012; Magee & Grienberger, 2020). Yet, because directly measuring both neuronal activity and synaptic changes in-vivo on large populations of neurons remains a technical challenge, little is known of the plasticity rules at play during behavior (Aljadeff et al., 2021; Graupner et al., 2016; Lim et al., 2015).

Even in the hippocampus, a brain area essential for memory (Morris, 2006) and where synaptic plasticity has been investigated most intensively (Bliss et al., 2018; Buchanan & Mellor, 2010), the learning rules that shape neuronal representations are only starting to be understood. For example, during familiarization to an environment by repeated unidirectional exploration, spatial representations in the CA1 subfield of the hippocampus gradually drift backwards (Dong et al., 2021; I. Lee et al., 2004; Mehta et al., 1997, 2000; Priestley et al., 2022; Roth et al., 2012) and this neural correlate of incidental learning is known to be dependent on the molecular machinery for LTP (Burke et al., 2008; Ekstrom et al., 2001; Kaganovsky et al., 2022). The population backward drift is faster in novel environments, slows down with familiarization, and occurs to a lesser degree in CA3, the main source of inputs to CA1 (Dong et al., 2021; Roth et al., 2012). Overall, this form of representational drift, resulting from shifts in the position of individual place fields (PFs), is thought to reflect ongoing synaptic plasticity. However, the precise rules and mechanisms explaining differences between familiarity levels and hippocampal subfields are unknown.

A fruitful approach to uncover the synaptic mechanisms supporting cognition has been to use computational modeling to infer the rules that would best fit in-vivo recordings (Aljadeff et al., 2021; Milstein et al., 2021). Early computational models suggested that classic Hebbian spike-timing-dependent-plasticity (STDP) (Bi & Poo, 2001) could cause individual PFs to shift backwards (Mehta et al., 2000; X. Yu et al., 2006). The mechanism is intuitive: the asymmetry of the rule favors potentiation of inputs that fire before the output place cell and depress inputs that fire after, such that, combined with repeated unidirectional track traversals, the output cell fires earlier on the track. However, these models were proof-of-concepts that used parameters potentially inflating the effects of STDP without systematically exploring the parameter space. As such, they do not account for the diversity in the dynamics of single PFs, which do not all shift backward and can occasionally shift forward, nor do they explain differences between hippocampal subfields and familiarity levels (Dong et al., 2021). Moreover, the effect of classic STDP was not compared to other phenomenological rules. Indeed, classic STDP is an imperfect way to describe synaptic plasticity. First, the STDP kernel itself can vary in shape and amplitude at CA3-CA1 synapses, depending on induction protocols (Inglebert et al., 2020; Wittenberg & Wang, 2006). Furthermore, Hebbian STDP rules in general have been undermined because 1) their impact may be too weak in natural regimes of firing and physiological conditions (Graupner et al., 2016; Inglebert et al., 2020; Lisman & Spruston, 2010) and 2) they operate on timescales too short to support the association of stimuli presented seconds apart (Gallistel & Matzel, 2013).

A promising alternative to explain PF dynamics could be behavioral-timescale-synaptic-plasticity (BTSP), a new type of non-Hebbian plasticity recently discovered at the CA3-CA1 pyramidal synapse (Bittner et al., 2017; Fan et al., 2023; Magee & Grienberger, 2020; Milstein et al., 2021). BTSP has three main differences with STDP: 1) it is triggered by rare but large dendritic calcium plateau potentials generally accompanied by a somatic burst of activity called a complex spike, 2) the induced synaptic changes are larger, 3) it operates on the timescale of seconds. The phenomenon originally discovered was a purely potentiating rule (Bittner et al., 2017), but the amplitude and polarity (potentiation vs depression) may be weight-dependent (Milstein et al., 2021) or depend on interactions with additional heterosynaptic rules with homeostatic effects (Chistiakova et al., 2015). So far, BTSP has mostly been considered as a mechanism underlying PF emergence (Fan et al., 2023; Magee & Grienberger, 2020; Priestley et al., 2022) or remapping (Milstein et al., 2021). Yet, because dendritic plateaus can spontaneously occur in neurons with an already established PF (Bittner et al., 2015; Cohen et al., 2017; Fan et al., 2023) and cause a PF translocation (Milstein et al., 2021), we hypothesized that a series of BTSP-triggering plateaus during exploration could lead to a PF shifting backward or forward, depending on the probability and location of such events.

Here, we used computational modeling to test the effect of different STDP and BTSP rules on PF shifting and compared our simulations to experimental observations from large populations of CA1 and CA3 neurons (Dong et al., 2021). The large sample-size afforded by 2-photon calcium imaging allowed us to accurately assess the variability in the shifting dynamics of single PFs. From this, we inferred that BTSP is more likely than STDP to support the evolution of hippocampal representations during learning, we deduced differences in the phenomenology of BTSP between CA1 and CA3 and we determined the dynamics of BTSP-triggering events as a function of familiarity.

## RESULTS

To assess the synaptic plasticity rules at play in the hippocampus during familiarization to new experiences, we used our previously published dataset of CA1 and CA3 pyramidal cells recorded in wild-type mice with 2-photon calcium-imaging (Dong et al., 2021). 11 animals (4 for CA1, 7 for CA3) were recorded while unidirectionally running multiple laps through a virtual linear track in a familiar environment and then switched to a novel virtual environment (Methods). We considered the lap-by-lap dynamics of the center-of-mass (COM) of individual PFs (2235 in CA1, 414 in CA3) as a proxy for ongoing reorganization of their synaptic weights. Our approach was 1) to characterize PF COM dynamics, and the differences between hippocampal subfields and familiarity levels, and 2) to model different plasticity rules and explore their parameter space to match the experimental data and infer the mechanisms that control different aspects of PF dynamics.

As reported in Dong et al. (2021), we found that many PFs are stable from lap-to-lap whereas some seem to linearly shift their position, usually backward but occasionally forward (Fig 1A). Here, we quantified shifting dynamics by performing a linear regression on the COM trajectory of each PF (Fig 1A-B). For all experimental conditions, there was a sizeable proportion of significantly shifting PFs, spanning a large range of shifting speeds. There were also clear differences between hippocampal subfields and familiarity levels (Fig 1B-D). CA3 had a lower proportion of shifting place fields, and shifts were slower than in CA1, particularly in the novel environment. Familiarity had a strong effect on the proportion of shifting place fields, with a large decrease in backward shifting PFs in familiar contexts for both CA1 and CA3. In addition, we noticed in CA3 an increase in the fraction of forward shifting PFs in the familiar context, making it higher than in CA1 (Fig 1B, C). These effects were visible in the full population of PFs and consistent across all mice.

**Figure 1.**
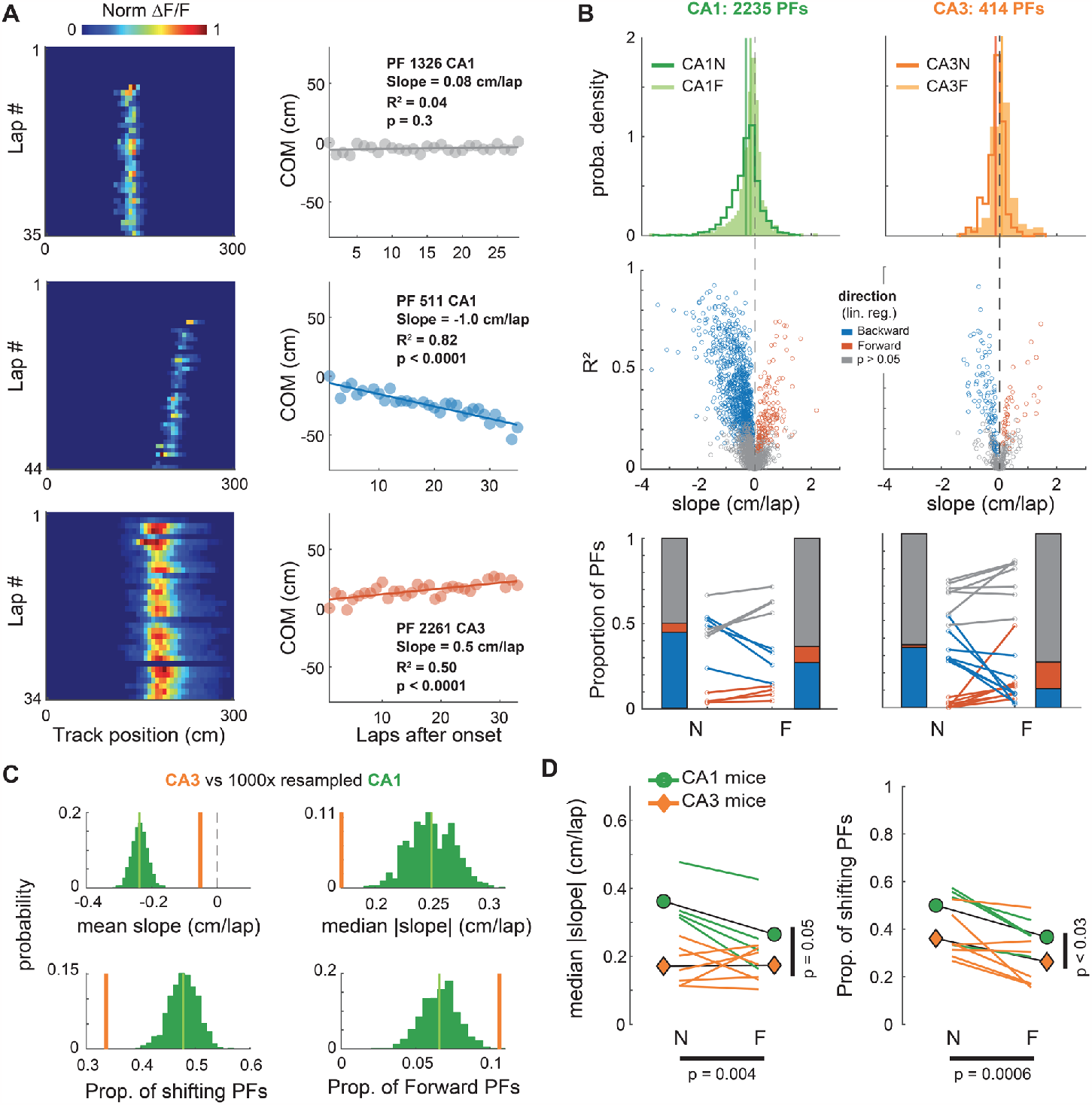
Linear shifting of place-fields decreases with familiarity and differs between CA1 and CA3. **A**.*Left*: Examples of CA1 and CA3 place-fields (PFs) recorded using 2-photon calcium imaging. *Right*: Linear regression on the onset-centered PF Center of Mass (COM) was used to classify each PF as shifting backward (blue), forward (red) or not significantly shifting (grey). **B**.Characterization of PF linear shifting in CA1 (left) and CA3 (right) for PFs defined over a span of at least 15 laps. *Top*: Probability density distributions of slopes (i.e. shifting speeds) for all PFs in CA1 and CA3 during navigation along a novel (N) or familiar (F) virtual track (CA1N: 1167 PFs, CA1F: 1068 PFs, CA3N: 235 PFs, CA3F: 179 PFs). *Middle*: estimated shifting speed vs linear regression fit (R^2^) for individual PFs. *Bottom*: Lines correspond to individual mice, stacked bars to averages across mice (n = 4 for CA1, 7 for CA3. Paired t-tests: CA1 N vs F backward: t(3) = -4.5, p = 0.020, forward: t(3) = 3, p = 0.057; CA3 N vs F backward: t(6) = -3.9, p = 0.008, forward: t(6) = 2.4, p = 0.051). **C**.Resampling exact tests controlling for the sample size difference between CA1 and CA3. 414 of the 2235 CA1 PFs were randomly resampled 1000 times to match CA3. The CA3 value was outside the resampled distribution for all statistics (green distribution). **D**.Animal-wise statistical tests (colored lines are individual mice, symbols are averages across mice). ANOVAs with repeated measures based on linear mixed effects models show effects of both the recorded subfield (CA1 vs CA3) and the environment familiarity (N vs F) on the proportion and speed of PF shifting. *Left*: Median Absolute Slope ∼ 1 + Subfield * Familiarity + (1 + Familiarity | Mice): Subfield: F(1,18) = 4.27, p = 0.053; Familiarity: F(1,18) = 11.14, p = 0.0037; Interaction: F(1,18) = 7.5, p = 0.013. Because the interaction was significant, we performed post-hoc paired t-tests with Bonferroni corrections for 4 comparisons: CA1N vs CA3N p = 0.0047, CA1N vs CA1F p = 0.078, CA1F vs CA3F p = 0.37, CA3N vs CA3F p = 1. *Right*: Proportion of shifting PFs ∼ 1 + Subfield + Familiarity + (1 + Familiarity | Mice): Subfield: F(1,19) = 5.59, p = 0.029; Familiarity: F(1,19) = 16.77, p = 0.0006; The interaction was excluded because it was not significant.

### STDP is too weak to explain PF shifting dynamics

Past computational studies suggested that backward shifting in CA1 could result from STDP at synapses from spatially modulated inputs (D’Albis et al., 2015; Mehta et al., 2000; X. Yu et al., 2006). We thus sought to determine whether the distribution of PF shifts that we observed could be explained by such classic Hebbian plasticity. We designed a simple model of a spiking place cell with stochastic and plastic inputs following a classic STDP rule (Fig 2A-B, Methods). Our model is inspired by seminal studies (Mehta et al., 2000; X. Yu et al., 2006) but differs from them in several ways (Table 2, Methods). Importantly, input parameters were adjusted to ensure an output with firing rates and PF widths as measured from CA1 recordings in mice (Fig S1, 2C). In contrast to past reports, this model produced few significantly backward shifting PFs, with a narrow range of small shifting speeds (Fig 2C) unlike what we observed experimentally in CA1 (Fig 1B, 2E). Past models used higher firing rates, which leads to a higher number of pre-post spike pairs and consequently increases the impact of STDP. We thus varied input parameters (Fig 2Fi) to cover a wide range of realistic and unrealistic output firing rates (Fig 2Fii and S1D — unrealistic Peak FR > 32Hz). Consistent backward shifting as reported by previous models occurred only for highly unrealistic output firing rates (Fig 2D-F). However, this increase in firing rate was not able to produce a range of shifting speeds as large as in our recordings, with shifting being exclusively backward and shifting speeds still constrained to small values (Fig 2Diii, E top, Fii). We explored the parameter space of our model extensively, but no set of parameters offered a good match to the data (Fig S2-7). Using CA3-like dynamic input PFs rather than static ones (Fig S2) improved the proportion of shifting PFs, but still yielded only small shifting speeds (Fig 2E, S5). Increasing the effect of STDP by allowing runaway potentiation and making the model more realistic by adding spike-rate adaptation to the output neuron and adding a dynamic delay in the update of synaptic weights did not improve the fit (Fig S3). Increasing the animal speed, which also amplifies the effect of STDP on backward shifting (because of the unidirectional movement) did not alter our conclusions either (Fig S5-7). Finally, because the exact amplitude and timescale of synaptic weight changes due to STDP protocols is not clear (Bi & Poo, 2001; Froemke et al., 2006; Mehta et al., 2000; Morrison et al., 2008; Shouval et al., 2010; Song et al., 2000; Wittenberg & Wang, 2006), we tested different combinations of parameters for the STDP rule (Fig S6-7). Realistic variations in the amplitude of weight changes and the time constants did not change our results; only unrealistically high values yielded consistent backward shifting (Fig S6-7B, C), without ever matching the range of shifting speed of our experimental data. Overall, the main effect of STDP is not PF backward shifting, which is weak, but it is an increase in output firing rates leading to PF enlargement (Fig S3D, S4, S6-7G-H, J-K). This PF width increase is not apparent in our recordings (Fig S4I-J), providing additional evidence that classic STDP is unlikely to be the mechanism underlying PF shifting dynamics in the hippocampus.

**Figure 2.**
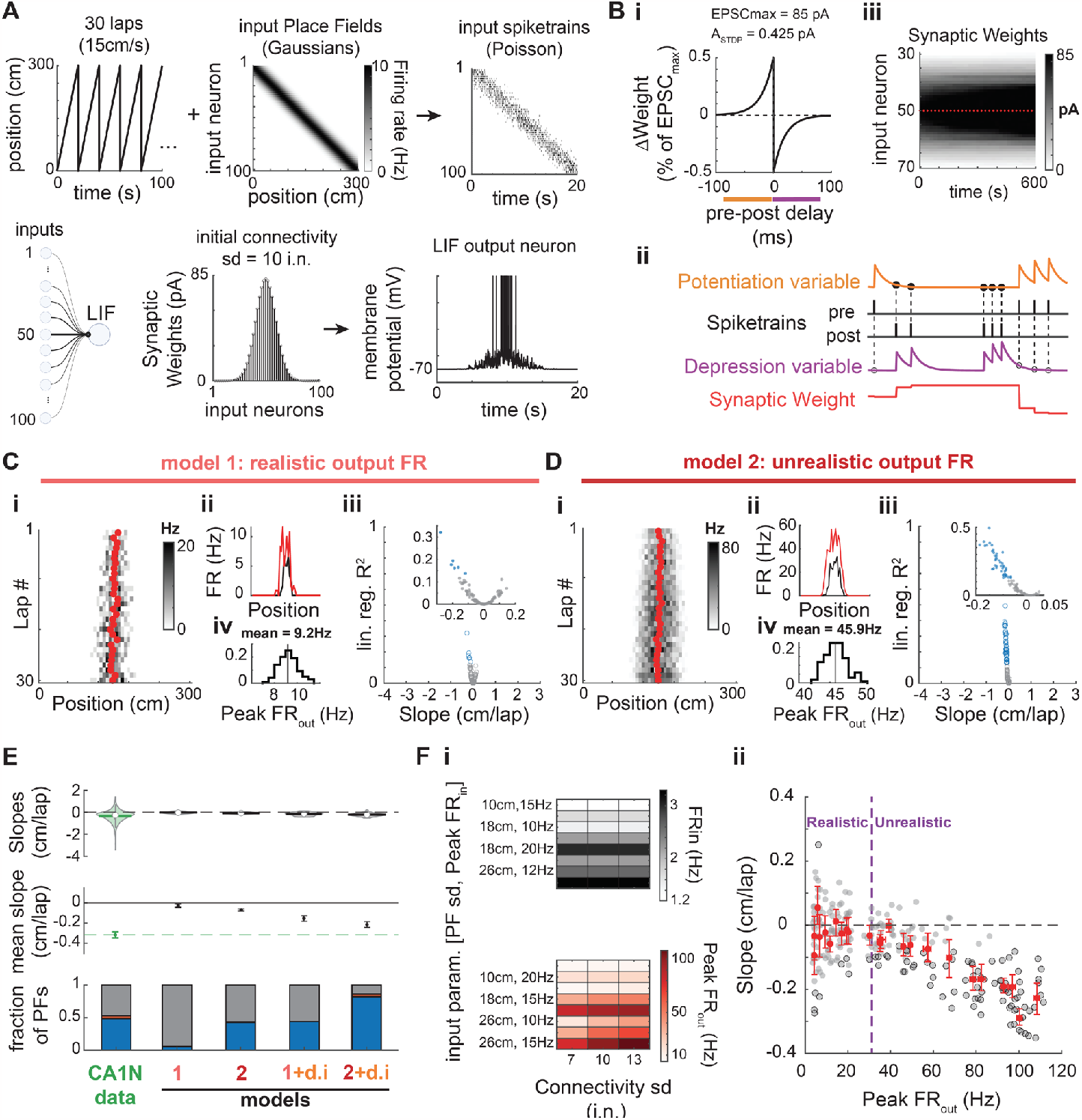
STDP does not explain PF shifting in CA1. A.Place Field model. A virtual animal runs at constant speed for 30 unidirectional laps (only 5 laps shown in upper left). Connectivity standard deviation (sd) expressed in number of input neurons (i.n.). **B.i**: The synaptic weight of each input neuron is updated at every time step (1 ms) according to a classic anti-symmetric STDP rule (20 ms time constant, maximum weight change A_STDP_ = 0.5% of EPSCmax = 0.425 pA). Synapses saturate at 85 pA. **ii**: The STDP rule is implemented via two plasticity variables triggered on pre or post spike-trains and used for updates at the time of each post or pre spike, respectively (see methods). **iii**: Evolution of the synaptic weights during an example 30-lap simulation. Red plus-signs mark the start of a new lap. **A-B**. Parameter values noted here correspond to the baseline model (model 1, panel C). **C**.Baseline model. **i-ii**: Example simulation that resulted in a significantly shifting PF. **i**: Red dots are the lapwise center of mass. Compare with Fig 1A. **ii**: Firing rates averaged over the first 3 laps (black) and the last 3 laps (red). There is a modest increase in FR and PF width resulting in a slight backward shift. **iii-iv**: Simulation of 100 PFs with the baseline parameters. **iii**: Linear regression fit (R^2^) as a function of the shifting speed (slope of the regression) for 100 simulated PFs. Same color code as in Fig. 1B: only a few PFs show a weak (small shift and R^2^) but significant backward shift (blue data points). No significant forward shifting. **iv**: Distribution of the peak firing rate of the output PF (30-lap average) for all 100 simulations: output firing rates are realistic (see Fig S1). Y-axis: fraction of PFs. **D**.Same as in C except the Peak FR_in_ parameter was raised from 10 to 15 Hz. A large proportion of the 100 simulated PFs significantly but weakly shifted backward (iii). However, the high output firing rates (Peak FR_out_ ∼ 45.9Hz) are outside the normal range of CA1 PFs (i, iv). **i-ii**: Example of the simulated PF with the largest backward shift in panel iii. **E**.Comparison of the shifts measured in CA1 data during navigation of a novel environment (same data as in Fig 1) and four different models (100 simulations each): model 1 (baseline parameters, data in panel C), model 2 with higher input and output firing rates (data in panel D), a modified model 1 with CA3N-like dynamic inputs (d.i.) following the probability distribution of slopes shown in Fig. 1B-top, and a modified model 2 with CA3N-like dynamic inputs. **Top**: Violin plots of the slope distributions (median is open circle, mean is solid line). **Middle**: Bootstrapped mean slope and 95% confidence intervals of the distributions shown above. The 3 later models result in consistent backward shifting (significantly below zero) but not as large as in CA1 (green, dashed line). **Bottom**: Proportions of PFs with backward (blue), forward (red) and non-significant shifting dynamics (grey). Model 2 and 1+d.i. have a proportion of backward shifting PFs similar to CA1N (PFs from all animals pooled), but no forward shifting. Model 2+d.i. inherits some forward shifting from the CA3-like dynamic inputs but proportions do not match CA1. **F**.Effect of firing rates on PF shifting induced by classic STDP (see also Fig S5). **i**: A set of 3 parameters controlling the inputs and thus the output firing rates without changing the plasticity rule were systematically varied to test 24 different conditions. All other parameters were as in A-B. Input PFs were static. **Top**: FRin is an estimate of the mean firing rate of input neurons across the whole track. **Bottom**: Peak FRout averaged across 20 simulations for each condition (x-axis values of red dots in panel ii). **ii**: 20 PFs were simulated per conditions (grey dots) with significant shifts marked by a black edge. Red dots are the means for each condition, with bootstrapped 95% CI in the x and y-axes. Dashed blue vertical line marks the upper bound of peak rates observed in CA1 in mice (Mou et al. 2018, see Fig S1): Consistent but modest backward shifting (without forward shifting) only occurs for unrealistically high output firing rates.

### BTSP explains PF shifting dynamics in CA1 and CA3

We next tested whether BTSP could support PF shifting dynamics by designing a new BTSP model that could easily replace the STDP rule in our initial spiking place cell model (Fig 3A-B). In contrast to past models that considered BTSP as a bidirectional plasticity rule (Cone & Shouval, 2021; Milstein et al., 2021), our strategy was to combine a pure potentiation rule as discovered by Bittner and colleagues (2017) with a simple homeostatic rule preventing runaway potentiation and maintaining the existence of a PF (i.e. not firing everywhere on the track) as observed in recordings from Milstein et al. (2021). The parameters of our model were optimized to fit Bittner et al. (2017) in-vitro experiments (Fig S8) and Milstein et al. in-vivo results (Fig S9-11 and Fig 3C). Our simulations of ‘Milstein-type’ PF translocation experiments revealed that combining a potentiation rule with homeostatic plasticity can lead to an apparent weight-dependent bidirectional plasticity rule (Fig S10-11) and is appropriate to study the effects of BTSP on PF shifting dynamics (see Methods). Overall, the advantage of our model is its simplicity, which allows: 1) to fit the Milstein dataset with a single set of parameters, 2) to implement BTSP in a network of spiking neurons, and 3) an easy comparison with the parameters of our STDP model from which it was adapted.

**Figure 3.**
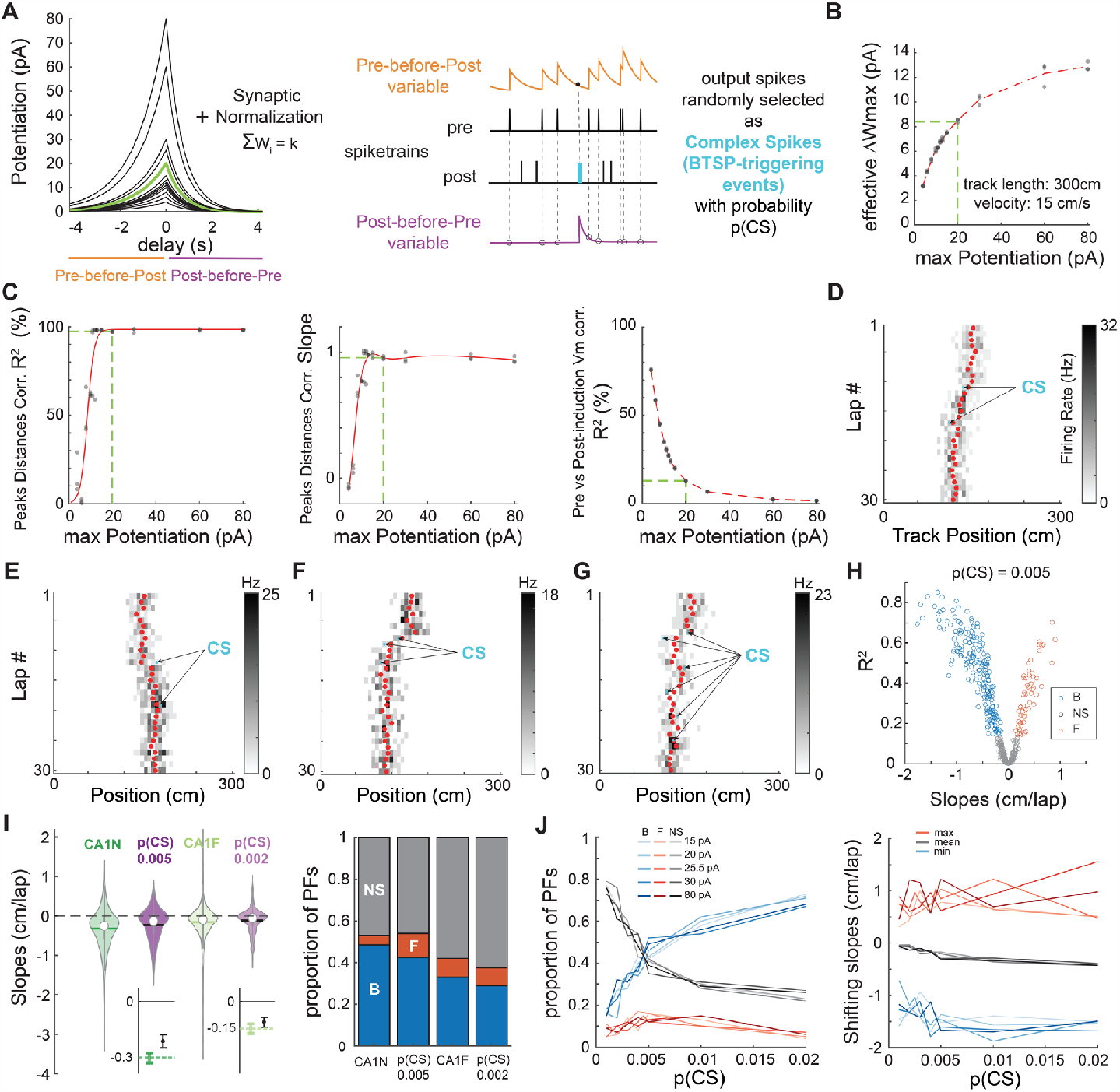
BTSP explains PF shifting in CA1. **A**.*Left:* Plasticity rules tested in B-C. The green kernel corresponds to the BTSP rule used in D-J. *Right*: Implementation of plasticity adapted from our STDP model. **B-C**. To determine a plausible maximum potentiation parameter for the BTSP rule, we tested different amplitudes (shown in A) in “Milstein-type” in-silico experiments as described in Fig. S11 (3 experiments per condition). **B**.Effective maximum weight change resulting from the combination of homeostatic plasticity and each potentiation rule in A. Estimated as in Fig S10H. The green dashed line corresponds to the green kernel in A. **C**.Optimization of the BTSP maximum potentiation parameter to fit Milstein et al. (2021)’s experimental findings (see Fig S10). The green dashed line indicates the optimal BTSP amplitude (minimal parameter value that maximizes the first 2 indicators and for which the third indicator is optimally low). **D-G**. Examples of 30-lap simulations of our place cell model (as in Fig. 2A) with plastic synapses following the optimized homeostatic BTSP rule (green kernel in A). Depending on the number and location of CSs (arrows), the COM trajectory (red dots) can go backward (**D, F**) or forward (**E**) and appear somewhat smooth and linear (**D**) or display abrupt shifts and changes of direction (**G**). **H**.Simulation of 500 PFs using p(CS) = 0.005. The distribution of backward, forward and non-significantly shifting PFs (assessed by linear regression of the COM as before) is reminiscent of CA1 (compare to Fig 1B). **I**.Comparison of the CA1 data (dark and light green, same as in Fig 1) with 2 versions of the model where only p(CS) was changed (dark and light purple). *Left*: Violin plots of the shifts distributions (median is open circle, mean is solid line). The models (500 simulated PFs each) cannot reproduce the most extreme shifts, but the variances are comparable to CA1. Insets on the bottom show bootstrapped means and 95% CI (small but significant difference between the model and CA1N, not significant for CA1F). *Right*: Proportions of backward (B, blue), forward (F, red) and non-significantly (NS, grey) shifting PFs. The models qualitatively match the data. **J**.Exploration of the parameter space: p(CS) and BTSP amplitude (maximum potentiation before synaptic normalization) were varied systematically. 100 simulations per condition. *Left*: Proportions of backward (blue), forward (red) and non-significantly (grey) shifting PFs. *Right*: minimum, mean and maximum shifts. The mean shift monotonically but only slightly decreases with p(CS) due to larger proportions of backward shifting PFs, not by inducing larger shifts.

To investigate the impact of BTSP on PF shifting during exploration, we simulated experiments as in Figure 2A (Fig 3D-J). Since the physiological causes of BTSP-triggering events (referred to in our model as “complex spikes”, or CSs, for convenience) are not well-understood (Magee & Grienberger, 2020), we considered that each output spike in the model had the potential to be a BTSP-triggering CS with a certain probability p(CS). Simulations using that strategy could lead to both backward and forward shifting PFs (Fig 3D-E). Because of the stochasticity in firing and in determining CSs, the model could produce smooth, sometimes linear-like trajectories (Fig 3D), but also yield more abrupt shifting when a CS occurred on the edge of the initial PF (Fig 3F), and even zigzag trajectories when multiple CSs occurred successively on different sides of the PF COM (Fig 3G). Large-scale simulations of 500 PFs with low p(CS) matched our experimental data well in terms of shifting speeds as well as proportion of shifting PFs (Fig 3H-I). By exploring the parameter space, we found that a familiarity-dependent decrease in p(CS) was sufficient to explain the lower amount of backward shifting in familiar environments (Fig 3I-J). Indeed, systematically varying p(CS) revealed that it directly controls the proportion of significantly shifting PFs but has little impact on the shifting speeds (Fig 3J), which is exactly the effect of familiarity in the experimental dataset (Fig 1). Testing different amplitudes for the BTSP rule, to control for edge effects in the effective maximum weight change (Fig 3B), did not alter our results (Fig 3J). Overall, we conclude that BTSP, unlike STDP, likely supports PF shifting dynamics in CA1 during familiarization.

Does BTSP also support PF shifting in CA3? A lower p(CS) than CA1 could potentially explain the smaller proportion of shifting PFs in CA3 (Fig 1B-D). However, forward shifting proportions are also different in CA3 than CA1 and Figure 3J shows that p(CS) or BTSP amplitude do not affect that proportion by much. As a result, the BTSP rule measured by Bittner et al. (2017) in CA1 could not fit our CA3 data well. We therefore hypothesized that a BTSP rule with different time constants could be at play in CA3. We found that, for a given p(CS), the extent of asymmetry of the BTSP rule strongly determines the ratio of backward/forward shifting PFs (Fig 4A). Because this ratio changes dramatically from familiar to novel environments in the experimental data, it suggests that the symmetry in the BTSP rule may be dynamic in CA3 (Fig 4B-D): in a familiar environment, the predicted BTSP rule must be close to symmetric, which is consistent with recent findings from in-vivo patch-clamp experiments (Li et al., 2023), whereas the very high ratio of backward/forward shifting observed during a novel experience is best explained by a highly asymmetric rule. Our simulations thus show that a BTSP rule different from CA1 could support PF shifting dynamics in CA3, with a familiarity-dependent change in its time constants, and lower p(CS) than in CA1.

**Figure 4.**
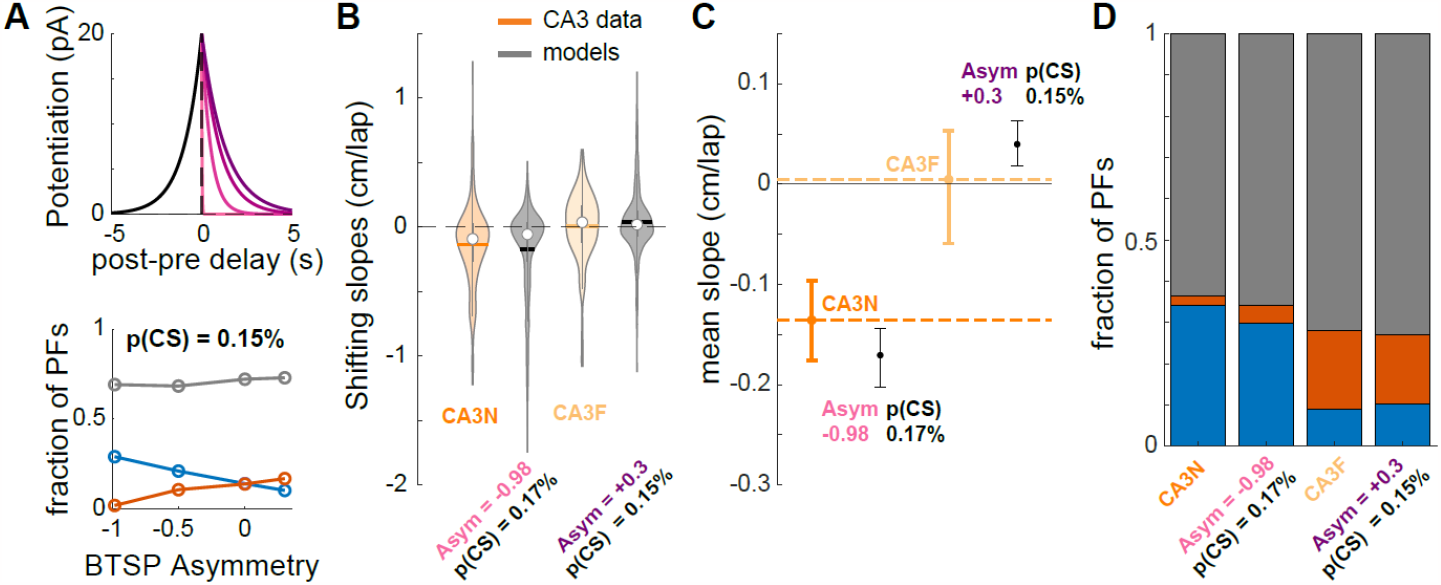
A dynamic BTSP rule supports PF shifting in CA3 in novel and familiar environments. **A**. *Top*. BTSP rules: τ_postpre_ was set at 1s, τ_prepost_ was varied from 20ms (lighter shade) to 1.3 s (darker shade). *Bottom*. Effect of BTSP rule asymmetry (τ_prepost -_ τ_postpre_) on the proportions of backward (blue), forward (red) and non-significantly shifting PFs (grey), when p(CS) is held constant (0.15%). 500 simulated PFs per condition. **B-D**. Comparison of the CA3 data (dark and light orange, same as in Fig 1) with 2 versions of the BTSP model (500 simulated PFs each). CA3N-like model parameters: p(CS) = 0.17%, τ_postpre_ = 1s, τ_prepost_ = 20ms. CA3F-like model parameters: p(CS) = 0.15%, τ_postpre_ = 1s, τ_prepost_ = 1.3s. **C**. Error bars are bootstrapped 95% CI of the mean.

### Nonlinear PF trajectories as signatures of the dynamic probability of BTSP-triggering events

If CS-triggered BTSP is the mechanism underlying PF shifting in the hippocampus, COM trajectories in experimental data should frequently look non-linear as they do in our simulations (Fig 3E-G). However, experimental reports have mostly focused on linear trajectories. We thus asked whether we could detect different types of COM trajectories in our experimental dataset (Fig 5-6). First, we used an unsupervised approach, performing principal component analysis (PCA) on the ensemble of COM trajectories from our CA1 and CA3 recordings (Fig 5A-B). This dimensionality reduction analysis revealed one main component explaining 77% of the variance in COM trajectories (Fig 5B, Fig S12). This template trajectory was non-linear with a large shift occurring through the first few laps. Further analysis, including all principal components, did not reveal meaningful clusters, suggesting that hippocampal PF shifting dynamics, regardless of familiarity levels, are best described as a continuum of a single type of non-linear plateauing trajectory but with different amplitudes and polarities (Fig 5C-D, S12B). Although different conditions did not define separate clusters, as confirmed by a separate non-linear dimensionality reduction method (Fig 5E), there were still differences in the distribution of shift amplitudes (absolute PC1 score) between conditions (Fig 5C), consistent with our initial linear regression analysis (Fig 1).

**Figure 5.**
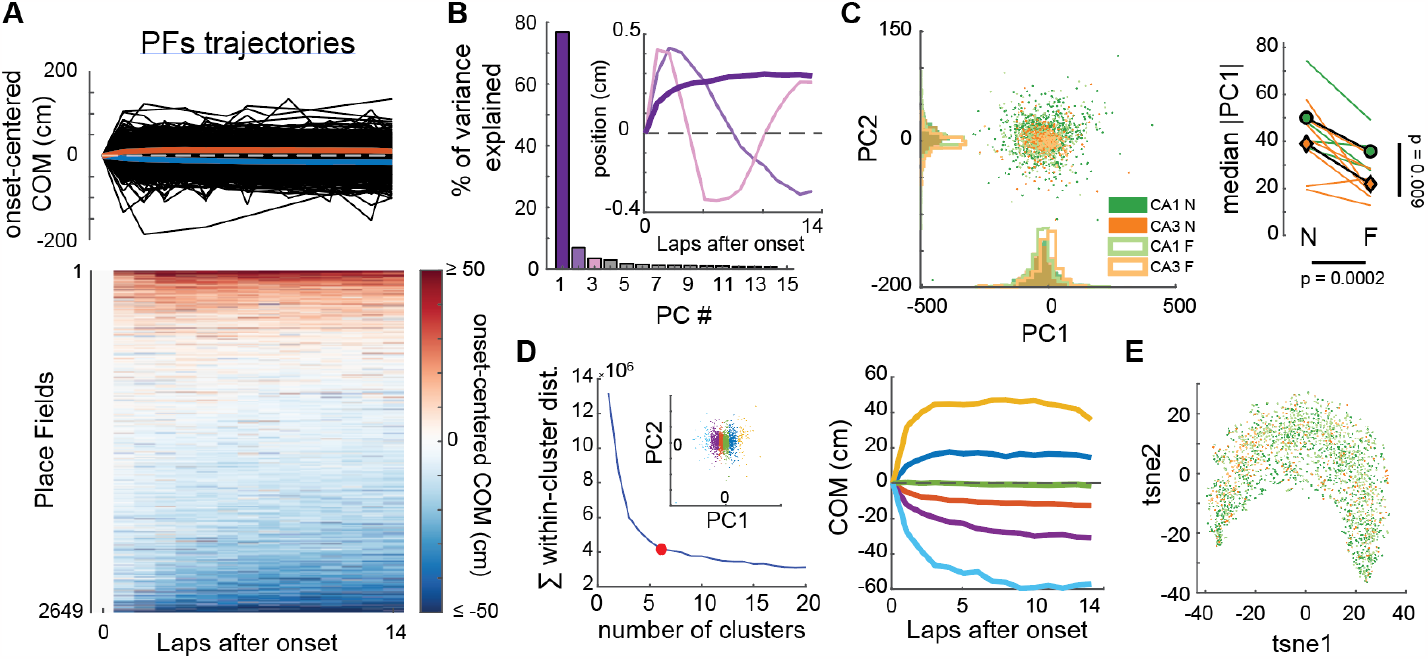
CA1 and CA3 PFs show a continuum of a single type of non-linear trajectory in experimental data. **A**.COM trajectories for all PFs recorded in CA1 and CA3 (same data as in Fig. 1). We used linear interpolation to infer the COM position on laps without activity, but results were similar without interpolation. *Top*: superimposed trajectories (black). Colored curves correspond to averages of PFs with negative (blue) or positive (red) average COM position. *Bottom*: same data in matrix form, each row being a PF. **B**.PCA was performed on the ensemble of trajectories shown in A. The first principal component PC1 explained 76.8% of the variance, revealing a non-linear trajectory template with a large shift during the first few laps (inset, dark purple bold curve — note that the polarity of the trajectory is irrelevant here because projection scores can be positive or negative, see Fig S12B). All other principal components revealed non-linearities but explained little variance each. **C**.*Left:* Scatterplot of the PC1 and PC2 projections of all recorded PFs, color-coded by subfield and familiarity. *Right:* Animal-wise ANOVA (see Fig.1D; colored lines are individual mice, symbols are averages). There is a significant effect of both the subfield and familiarity on PC1 scores. Median Absolute PC1 Score ∼ 1 + Subfield + Familiarity + (1 + Familiarity | Mice): Subfield: F(1,19) = 8.32, p = 0.0095; Familiarity: F(1,19) = 20.33, p = 0.00024; The interaction was excluded because not significant. **D**.*Left*: K-means clustering of all PFs trajectories using all principal components. Goodness-of-fit was optimal for 6 clusters (red dot = elbow), but clusters simply corresponded to segments of the PC1 scores (inset). *Right:* The color code is the same as in the left inset. Average COM trajectory for each k-means cluster reveals a continuum of the same PC1-like non-linear trajectory. **E**.non-linear dimensionality reduction (tSNE) confirmed the PCA analysis: COM trajectories do not form separate clusters but are spread along a continuum, with CA1N, CA1F, CA3N and CA3F PFs distributed in a salt-and-pepper fashion (color-code as in C).

**Figure 6.**
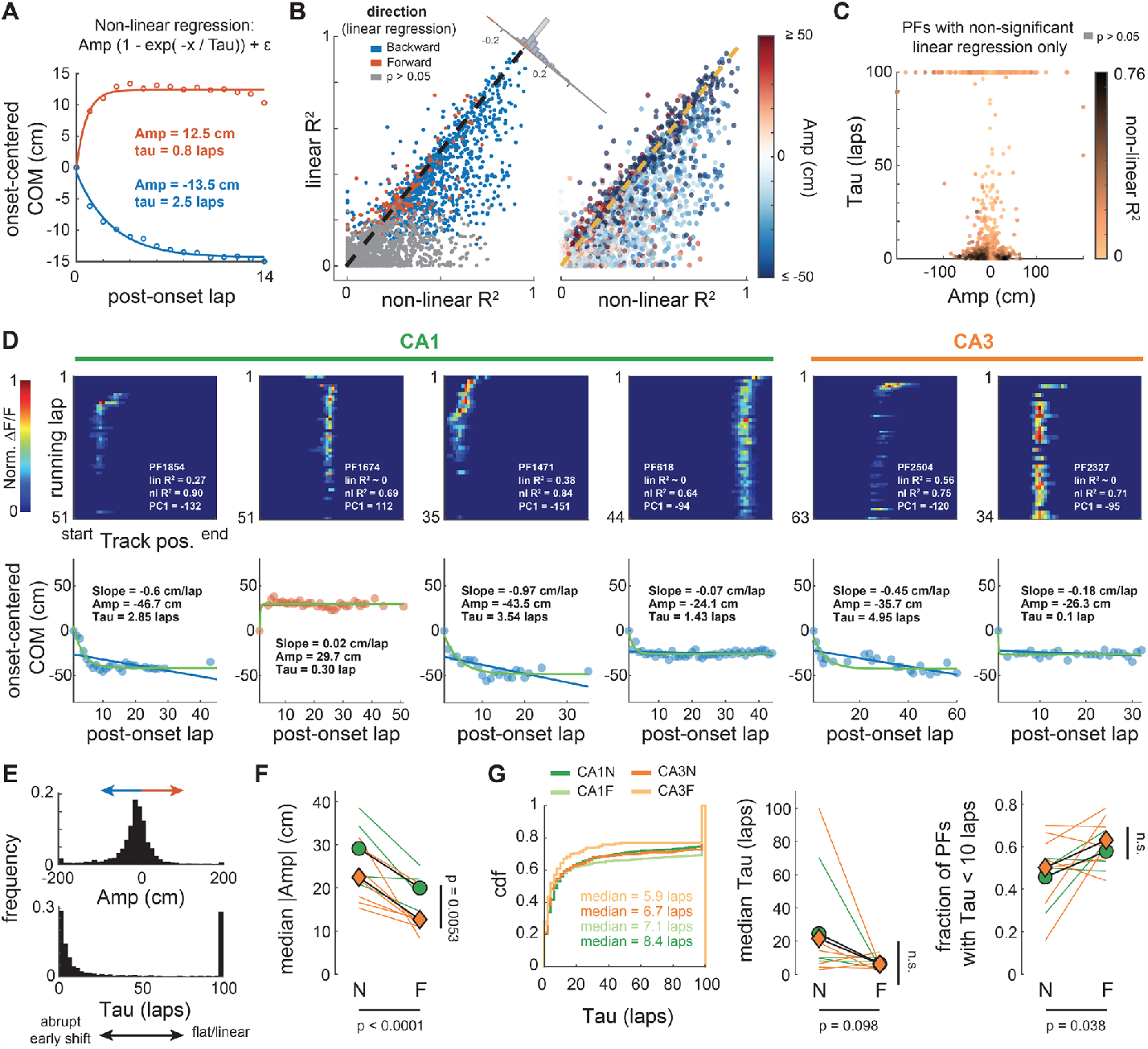
Single PF trajectories consistently show non-linear shifts early after PF emergence. **A**.Data points correspond to averages of PFs with negative (blue) or positive (red) mean COM position (as in Fig 5A). These averages are well fitted by a plateauing exponential function that captures features of PC1 in Fig. 5B. The function has 3 parameters: the amplitude Amp corresponding to the position of the plateau, the time constant Tau defining how fast the plateau is reached, and an intercept generally very close to 0. The sign of Amp describes the direction of the shift. The larger the Tau the flatter the trajectory, the shorter the more abrupt. **B**.Comparison of the goodness-of-fit (R^2^) between this non-linear regression and a linear regression (as in Fig 1) for all 2649 PFs. The non-linear regression fits most individual PF trajectories as well or better (data points under the identity dashed line). The difference of fit (corner histogram) is skewed towards positive values (Wilcoxon signed-rank test: z = 27.5, p < 0.0001), showing PFs are better described by a plateauing exponential (backward or forward shifting). **C**.Many PFs that were categorized as non-significantly shifting with the linear regression are well fitted (dark points) by a plateauing exponential with an abrupt shift (small tau), going backward or forward (Amp sign). **D**.Examples of PFs recorded in CA1 or CA3 with dynamics well described by a plateauing exponential. *Top*: lap-wise PF activity, with goodness-of-fit values (R^2^) for the linear and non-linear regressions, and the PC1 score for comparison. *Bottom*: linear (blue or red line) and non-linear (green curve) regressions on the lap-wise COM (data points). Backward shift (negative Amp) in blue, forward in red. Note that in some PFs the shift occurs on lap 1 after onset (e.g., PFs 618 and 2327) whereas in others the shift is more gradual. **E**.Distribution of Amp and Tau values for all PFs combined (see Amp and Tau covariance in Fig S13) **F**.Animal-wise ANOVA (see Fig.1D; colored lines are individual mice, symbols are averages) shows consistent effects of the subfield (CA1, green vs CA3, orange) and environment familiarity (N vs F) on the absolute amplitude. Median Absolute Amp ∼ 1 + Subfield + Familiarity + (1 + Familiarity | Mice): Subfield: F(1,19) = 9.88, p = 0.0053; Familiarity: F(1,19) = 25.97, p < 0.0001; The interaction was excluded because not significant. **G**.Cumulative density distributions pooling all PFs of a given condition (*left*) and animal-wise statistics (*middle, right*) for Tau. There is little difference between conditions in terms of Tau, with a modest but significant increase in the fraction of PFs with small Tau values (i.e. early shift) in familiar environments (*right*). *Middle*: Median Tau ∼ 1 + Subfield + Familiarity + (1 + Familiarity | Mice): Subfield: F(1,19) = 0.03, p = 0.86; Familiarity: F(1,19) = 3.02, p = 0.098. *Right*: Fraction of PFs with Tau < 10 laps ∼ 1 + Subfield + Familiarity + (1 + Familiarity | Mice): Subfield: F(1,19) = 0.92, p = 0.35; Familiarity: F(1,19) = 4.97, p = 0.038. Interactions were excluded because not significant.

To verify that individual PFs exhibited the type of trajectory identified by PCA, and to further characterize this phenomenon, we performed nonlinear regression and fitted a plateauing exponential to each COM trajectory (Fig 6). This supervised approach shows that most PFs are better described by this nonlinear trajectory than a continuous linear drift, for both backward and forward shifting PFs, allowing us to detect dynamic PFs previously considered not significantly shifting by the linear analysis (Fig 6B-D). The distribution of amplitudes and time constants reveals 3 classes of trajectories: early shifts, stable and linear-like (Fig S13), with a majority of PFs having an early shift (Fig 6E). These shifts can be very abrupt, occurring on the first lap after PF emergence, but they can also develop more slowly over the course of several laps (Fig 6D-E). Overall, the non-linear trajectories are consistent with a BTSP mechanism triggered by rare events. The prevalence of early shifts suggests that the probability of BTSP-triggering events is dynamic and that these events tend to occur soon after PF emergence. We checked for differences between conditions (Fig 3F-G, S14): in contrast to shift amplitudes, with less shifting in CA3 and familiar environments (consistent with previous analyses in Figures 1 and 5C), there was little evidence for differences in the dynamics of early shifts (Fig 6G). This suggests that the dynamics of p(CS) are similar across regions and familiarity levels.

The plateauing shape of the main component of PF trajectories shows that p(CS) is largest around PF emergence; but do BTSP-driven shifts occur later? The zigzagging shapes of the other components from the PCA (Fig 5), for which individual PFs can have a high score (Fig S12), suggest they do. Although later shifts seem rare, as evidenced by the lower amount of variance explained by these components, we found several examples of sinuous or zigzagging PF trajectories in our experimental data (Fig 7A) as predicted by our BTSP model (Fig 3G). Quantification of lap-to-lap COM displacement as a function of laps after PF emergence shows that these examples of zigzagging trajectories are representative of a global phenomenon: large shifts are more likely on the first laps but continue to occur with a constant, non-zero probability late after emergence (Fig S15, Fig 7B-C). Diffusion analysis, which considers the PF COM as a moving particle in a 1-dimensional space (Einstein, 1905), reveals that, after the first three laps, PF shifting dynamics follow a random walk with constant diffusion coefficient (Fig 7B-C). Comparison with computational models shows that such a random walk is not the product of stochastic firing but requires synaptic plasticity. In line with previous analyses (Fig 3-4), differences in the diffusion coefficient between familiarity levels and subfields can be explained by differences in p(CS).

**Figure 7.**
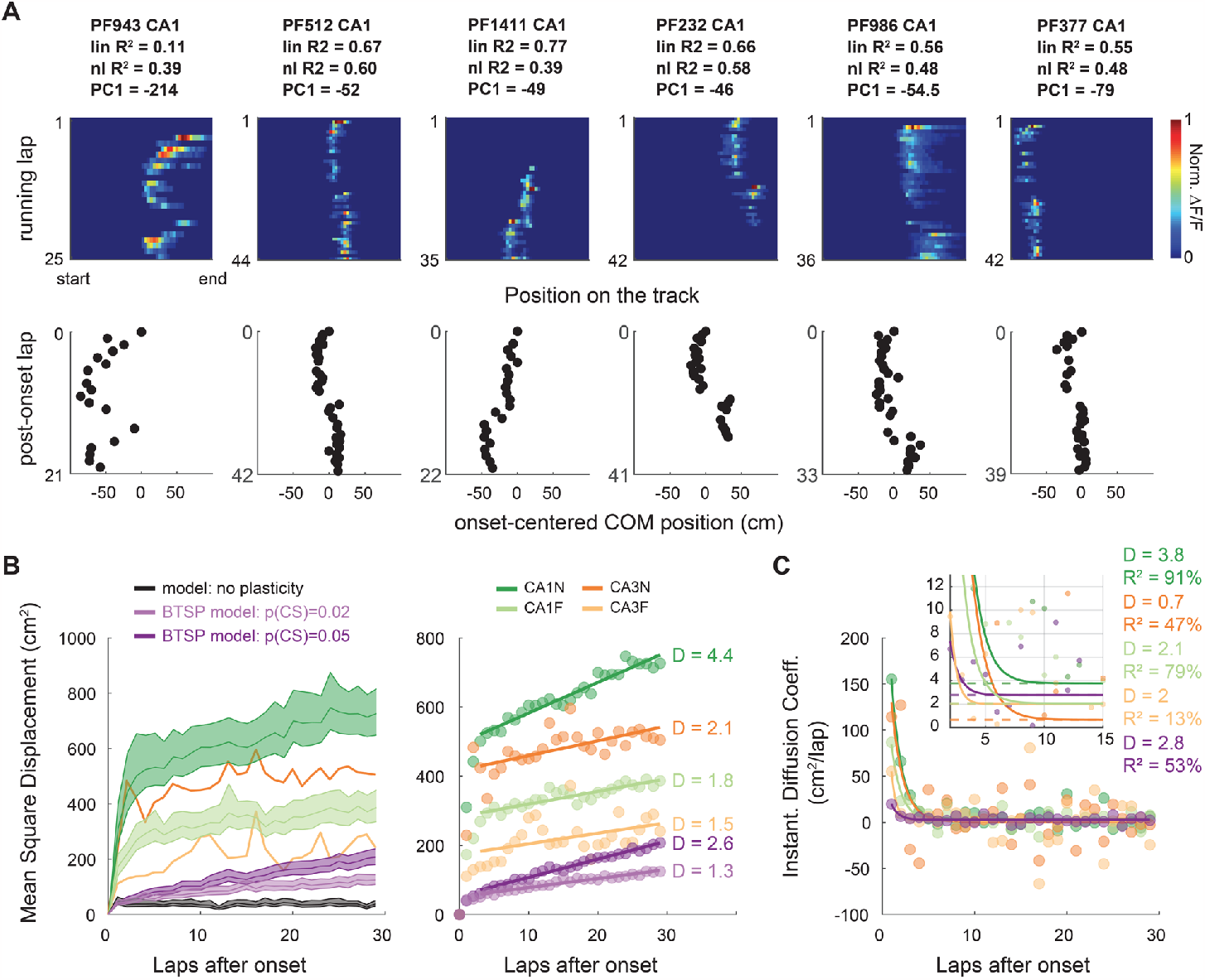
The probability of shift-inducing BTSP-triggering events decays to a constant after PF emergence. **A**.Example of PFs showing abrupt shifting late after PF emergence, resulting in zigzag COM trajectories (bottom). PF 1471 (CA1) and 2504 (CA3) in Fig. 6D are other examples with multiple shifting events. **B-C**. Diffusion analysis on PFs defined over at least 30 laps. **B**.Mean Squared Displacement of the COM (MSD) as a function of post-onset laps (computed over all PFs of each condition: n = 942 for CA1N, 880 for CA1F, 222 for CA3N,100 for CA3F and 500 for each model). For a random walk in a 1D environment such as our linear track, MSD = 2*D*laps, D being the diffusion coefficient. *Right*: MSD with 95% bootstrapped confidence interval for each condition and model. The large CIs of CA3 were omitted for readability. *Left*: linear regression on the MSD from lap 4 to lap 30 shows that PF shifting after lap 3 is well explained by a random walk with constant diffusion coefficient D. CA1N: R^2^ = 94.2%, p < 0.0001; CA1F: R^2^ = 80.1% p < 0.0001, CA3N R^2^ = 43.1% p = 0.0002, CA3F R^2^ = 20.1% p = 0.019, BTSP model p(CS) = 0.005: R^2^ = 98.3%, p < 0.0001; BTSP model p(CS) = 0.002: R^2^ = 94.5%, p < 0.0001. **C**.Alternative method of estimation of D by fitting the derivative of MSD (data points) to a decaying exponential p1*exp(-(x-1)/p2)+p3. The asymptotes (p1 parameter, dashed lines) correspond to D and qualitatively match the values estimated by linear regression in panel C.

## DISCUSSION

From our study emerges the view that: 1) BTSP rather than STDP supports the single-cell shifting dynamics of hippocampal representations during exploration of an environment, 2) the probability of BTSP-triggering is maximal at PF onset and then decays to a constant, thus driving a random walk of PFs after a few laps, 3) the probability of BTSP-triggering events is higher during novel experiences and 4) BTSP-triggering events also occur in CA3, with similar dynamics but a lower average probability and a different BTSP rule than CA1, switching from asymmetry to symmetry with familiarization. These BTSP-induced changes in spatial representations are a form of fast representational drift that could support continual learning during ongoing experience or help pattern separation to discriminate events close in time (Masset et al., 2022; Mau et al., 2020).

Our modeling suggests that the PF shifting dynamics induced by classic Hebbian STDP do not cover the range of trajectories observed in PFs recorded in the hippocampus (Fig 2). This conclusion contrasts with that of seminal studies (D’Albis et al., 2015; Mehta et al., 2000; X. Yu et al., 2006). We showed that this discrepancy comes from the fact that 1) previous models used unrealistically high firing rates, which enhances the effect of STDP by increasing the number of pre-post spike pairs, and 2) we had access to a larger sample of recordings to compare to. Using realistic firing rates, we found that STDP is too weak to induce the large shifting speeds that often occur in real PFs.

A counterargument could be that our model did not consider some complexities of real place cells. For instance, neurons of the hippocampal formation tend to fire at different phases of the theta rhythm, with CA3 inputs repeatedly firing before superficial CA1 place cells (Valero & De La Prida, 2018), which would amplify the potentiating effects of STDP and increase backward shifts. Similarly, phase precession in the CA3 inputs was shown to increase shifting speeds up to what we experimentally measured (D’Albis et al., 2015), although that effect may be dampened if precession in CA1 is not fully inherited from CA3 (and with the use of lower, more realistic firing rates). Overall, in the case of familiarization to a new environment, improving the realism of the place cell model by accounting for phase-preference and precession may at best amplify the linear backward shifting due to the asymmetry of the STDP rule; it cannot explain the higher-than-chance proportion of forward shifting (Fig 1-2) nor the nonlinear trajectories (Fig 6) that are more representative of the global phenomenon than a linear drift (Fig 5, 7). These aspects of PF dynamics were not characterized before, but they are consistent with previously reported examples of forward shifting PFs (I. Lee & Knierim, 2007; Roth et al., 2012) and PFs with nonlinear trajectories (Kaganovsky et al., 2022; I. Lee & Knierim, 2007; Priestley et al., 2022).

Could an improved model of classic synaptic plasticity, accounting for STDP but also for rate and heterosynaptic effects (Inglebert et al., 2020; Keck et al., 2017; Shouval et al., 2010; Zenke et al., 2015) better explain hippocampal PF shifting dynamics? Previous attempts suggest that it is not sufficient to yield large enough PF shifts (D’Albis et al., 2015; X. Yu et al., 2008). In contrast, our study shows that non-Hebbian BTSP is a clear way to explain hippocampal PF shifting because, unlike other known plasticity rules, it causes large synaptic weight changes and is triggered by rare dendritic events (associated to CSs) that can induce nonlinear shifts, both backward and forward depending on where on the track the CSs occur (Fig 3). In our model, the probability of BTSP-triggering events controls the proportion of significantly shifting PFs (Fig 3J) and the asymmetry of the BTSP rule determines the ratio of backward vs forward shifting PFs (Fig 4). Surprisingly, varying the amplitude of BTSP did not strongly affect the magnitude of shifts (Fig 3J), at least not in the range that we investigated, but this is due to dampening effects of the simple homeostatic rule that we used. In theory, the magnitude of shifts should depend on three factors: the location of BTSP-triggering events, the amplitude of BTSP and its time constants (the larger the time-constant, the more inputs are potentiated).

BTSP is a recent discovery and its phenomenology and mechanisms are not fully worked out. The original finding suggested a purely potentiating rule (Bittner et al., 2017), which would lead to runaway potentiation. Even with bounded synaptic weights, this rule would eventually saturate all synapses, in contrast to what recent experiments showed: two successive BTSP-triggering events potentiated inputs near the second CS location but depressed activity at other locations (Milstein et al., 2021). Recent work has suggested that a weight-dependent bidirectional homosynaptic rule could explain the phenomenon (Cone & Shouval, 2021; Milstein et al., 2021) but these models did not consider alternatives involving interactions between the original BTSP potentiating rule and fast heterosynaptic effects known to prevent runaway synaptic dynamics (Chistiakova et al., 2015; Zenke & Gerstner, 2017). Heterosynaptic competition and cooperativity are prevalent in the hippocampus (Chater & Goda, 2021; Magó et al., 2020) and can modulate BTSP (O’Dell, 2022). To model BTSP, we thus chose to combine the original BTSP rule with synaptic normalization, a simple heterosynaptic rule mediating homeostasis. The simplicity of that strategy allowed us to implement BTSP in a spiking network, optimize a single set of parameters to match the most recent experimental data (Fig S10) and was important to directly compare with our results on STDP. However, synaptic normalization is not realistic and induces some limitations in our model (see Methods). Therefore, to determine whether heterosynaptic or purely homosynaptic processes support the bidirectional changes observed in BTSP-induction experiments, future comparisons between the two classes of models should implement more realistic fast heterosynaptic rules (Abraham, 2008; Chistiakova et al., 2015; Ebner et al., 2019; Moldwin et al., 2023; Triesch et al., 2018). Additionally, experiments with longer tracks and varying speeds will be required to rigorously test each model’s predictions on the effect of BTSP-triggering events occurring more than 5s away from the initial PF.

Regardless of the homo- or heterosynaptic nature of BTSP, our study identifies several phenomenological aspects of BTSP.

First, our simulations of BTSP-induction experiments precisely quantified the amplitude of synaptic weight changes with BTSP (Fig S8-10): the maximum weight change due to a input spike-CS pairing was ∼4 pA in single-input in-vitro stimulations (Bittner et al., 2017) and 6-8 pA in PF translocation experiments (Milstein et al., 2021), that is 8 to 16 times higher than for STDP (∼0.5pA).

Second, our modeling of spontaneous PF dynamics during exploration, in combination with our characterization of PF trajectories in-vivo, provides crucial information on when and how often BTSP-triggering events occur. First, we found that these events are most likely at or right after PF onset, often leading to abrupt early shifts (Fig 5), which is consistent with the idea that BTSP-triggering dendritic plateaus is a major mechanism underlying PF emergence (Bittner et al., 2017; Fan et al., 2023; Priestley et al., 2022; Sheffield et al., 2017). Second, we extend previous in-vivo research that reported CSs in CA1 neurons with an already established PF (Bittner et al., 2015; Cohen et al., 2017; Fan et al., 2023; Milstein et al., 2021) by providing evidence that BTSP-triggering events do happen long after PF formation, with a dynamic probability that relaxes a few laps after PF emergence to a non-zero constant (Fig 7). Finally, since our model, which assumes that CSs occur in-field, does not explain the largest shifts observed in CA1 (Fig 3, 7), it suggests that BTSP-triggering events occasionally happen out-of-field.

Interestingly, direct measures of the frequency of CSs are inconsistent across studies, likely due to low sample sizes (7 to 30 cells). Bittner et al. (2015) found an average of 1.8 CSs per 100 spikes in a familiar environment, with higher p(CS) during the peak of theta-oscillations, whereas Cohen et al. (2017) reported much lower frequencies. Fan et al. (2023), using voltage imaging rather than patch-clamp, detected many CSs of short duration. Our analysis, based on hundreds of PFs, points to a very low probability of BTSP-triggering events (∼0.2 per hundred spikes in a familiar environment, which is an upper bound, given that some shifting may be inherited from dynamic CA3 inputs). Our results thus suggests that not all experimentally recorded CSs necessarily trigger BTSP, or not to the same degree, perhaps depending on the duration of the dendritic plateau potential (Takahashi & Magee, 2009). Our finding that the probability of BTSP-triggering event decays after PF emergence could thus be due to shorter CSs in established place cells (Bittner et al., 2015; Fan et al., 2023).

Finally, our study suggests that BTSP is not restricted to CA1, where it was discovered, but also occurs in CA3 in vivo, albeit with phenomenological differences (Fig 4). Dendritic calcium plateaus and associated CSs are indeed not specific to CA1; they have been recorded in cortical (Xu et al., 2012) and CA3 pyramidal cells (Balind et al., 2019), but their role in plasticity and their probability of occurrence was not known. Our study suggests that, in CA3, they can trigger BTSP, inducing PF shifting, and that their probability follows similar dynamics as in CA1, decaying after PF emergence (Fig 5-7). Unlike CA1 however, we found that: 1) the lower proportion of shifting PFs demonstrates a lower probability of BTSP-triggering events, even in new environments, 2) the smaller shifts suggests that fewer events occur out-of-field, and 3) the BTSP rule must be close to symmetric to explain the equal proportion of forward and backward shifting in familiar environments. This third point is consistent with recent in-vivo patch-clamp experiments that measured symmetric potentiation profiles following spontaneous and induced CSs (Li et al., 2023). Intriguingly, in new environments PF shifting proportions are dramatically skewed backwards, suggesting a highly asymmetric rule. This novelty-dependent change in the time-constants of the BTSP kernel could be carried by changes in the duration of the dendritic plateaus: CSs appear longer in CA3 than CA1 in familiar environments (Li et al., 2023), but a CA3-specific short calcium spike (Magó et al., 2021) could be more prevalent in novel contexts.

To conclude, our study of PF shifting dynamics offers a unique perspective on the synaptic mechanisms at play during incidental learning and memory formation. It shows that 1) BTSP drives the dynamics of hippocampal representations during familiarization, 2) the average probability of BTSP-triggering events is higher during novel experiences than familiar ones, especially in CA1, and the shape of the BTSP rule changes with familiarity in CA3. Novelty-dependent neuromodulatory and inhibitory signals (Pedrosa & Clopath, 2020; Sheffield et al., 2017) could mediate these changes by modulating both synaptic eligibility traces and the probability and duration of BTSP-triggering calcium plateaus (Fuchsberger et al., 2022; Magee & Grienberger, 2020).

## METHODS

### Experimental recordings

All experimental data analyzed in this study were previously published in Dong et al., 2021. Experimental procedures were in accordance with the University of Chicago Animal Care and Use Committee guidelines.

Briefly, GCaMP6f was first expressed in CA1 or CA3 principal neurons of the dorsal hippocampus of 10-12 week-old male mice. AAV1-CamkII-GCaMP6f was injected in the CA1 subfield of C57Bl6 mice whereas, to specifically target CA3 and exclude other hippocampal subfields, a CRE-dependent version of the same genetically encoded calcium indicator was injected in Grik4-cre mice (C57Bl6 background). To record the activity of large populations of neurons, mice were head-fixed under the objective of a 2-photon microscope.

Mice behavior consisted of running on a Styrofoam wheel to move through a virtual environment displayed on surrounding screens. Mice could only go forward or backward on a 300 cm virtual linear track and were trained through positive reinforcement to run forward multiple laps. Water reward was provided at the end of each lap, at which point the display was paused for 1.5 s before the animal was “teleported” back to the start of the track to start a new lap. Five days after window-implantation surgery, mice were trained for 10-14 days in a virtual environment defined as the familiar context (F) until they reached a consistent speed criterion of at least 10 cm/s (i.e., more than 2 laps/min). Engagement with the virtual environment was evident from the preemptive licking and the slowing down shown by animals before they reached the reward zone at the end of a lap (Dong et al., 2021; Krishnan et al., 2022).

Imaging was performed on the following days: on day 1, mice were exposed to the familiar environment and allowed to run at least 20 laps, followed by an exposure to a new environment (N1) with another 300 cm track in which they were allowed to run in at least 35 laps. A similar procedure was followed on day 2, during which a new field of view was recorded, with mice navigating again the familiar environment and then switched to a second new environment (N2). In the present study, all place fields detected in day 1 and day 2 were combined and labelled either F or N.

## Data preprocessing

Motion correction of the raw movies, cell detection and signal extraction were performed as previously published using custom MATLAB scripts (Dong et al., 2021). Place fields (PFs) were identified and defined as in Dong et al. 2021 using a method combining criteria about the peak fluorescence, stability and size of the PF compared against chance levels (Dombeck et al., 2010; Dong et al., 2021; Grijseels et al., 2021; Sheffield et al., 2017). Note that the criteria previously established were loose enough to not exclude shifting PFs. PFs too close to the beginning or end of the track were excluded (Dong et al., 2021). PFs from the same cell were considered independent. All analyses were performed on PFs in which non-significant activity and activity outside the defined PF region was removed. PF onset, the emergence lap of a given PF, was detected as in Dong et al. 2021 by finding the first lap where 1) a significant transient occurred in the PF region and 2) this lap was followed by significant PF activity in 2 out of 5 of the following laps.

### Analysis of PF trajectories

Analysis of PF shifting dynamics was based on the center-of-mass (COM) of the lapwise binned activity of a given PF. The 300 cm track was divided in 50 spatial bins and the lapwise normalized fluorescence F_i_ was averaged for each bin i. The COM on lap n was computed as follows:

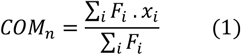

Where x_i_ is the position of bin i on the track (i.e. the distance from the start). For simulated data, the COM was computed in the same way except that F_i_ was the firing rate (i.e., the number of action potentials in bin i divided by the time spent in bin i). COM_n_ locations were then centered on COM_onset_, i.e., on the COM position during the emergence lap.

In recorded PFs, not all laps after onset necessarily show significant activity. For all analyses of the PF dynamics, we interpolated the COM location on post-onset laps without activity that were intercalated with active laps. If PF activity disappeared and did not come back during the session, the final laps without activity were excluded. For all analyses except in Fig 7B-C, we only included PFs that, after interpolation, were defined on at least 15 laps. For Fig 7B-C, the inclusion criterion was set at 30 laps or more (reducing the number of PFs but allowing a better picture of the long-term dynamics of PFs). An inclusion criterion of a minimum number of laps with a PF defined is an obvious but important prerequisite to assess PF dynamics. Moreover, interpolation and an inclusion criterion are necessary for the PCA and diffusion analyses (see below) and were thus implemented for all other analysis for comparison. However, neither the interpolation nor the minimum number of laps (0, 15 or 30) affected our conclusions.

To detect significant backward or forward linear shifts of the lapwise COM, linear regression between onset-centered COM position (response variable) and lap number n (predictor variable) was performed using the Matlab *regress* function.

For the non-linear regression analysis (Fig 5), we used the Matlab *fit* function with the nonlinear least square method and Trust-Region algorithm to fit the following function to each PF:

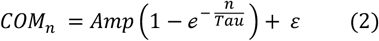

Where Amp (in cm), Tau (in laps) and ε (in cm) are the parameters to fit and n is the lap number. The parameter search starting point was [14 cm, 2 laps, 0 cm] if the linear regression slope was positive and [-15 cm, 2 laps, 0 cm] otherwise. Parameter search was bounded and stopped in case absolute(Amp) = 200 cm or Tau = 100 laps, or absolute(ε) = 25 cm.

To compare goodness-of-fit between linear and non-linear regressions, we chose to compare the respective R-squared statistics (coefficient of determination). We did not use the adjusted R-squared for the non-linear regression because the nonlinear model only has one parameter more than the linear model, overfitting was not a concern and the goal of the analysis was to determine the best description of a given PF trajectory, not to find an optimal model that would best predict out-of-sample data.

For the non-supervised analysis of PF trajectories (Fig 4), all PF trajectories (a PF trajectory being a vector of onset-centered COM position) were truncated to only include the first 15 laps where the PF was defined (i.e., 14 laps post-onset). We performed principal component analysis (*pca* Matlab function) on the matrix of all the truncated PF trajectories aligned on their respective onset lap using the singular value decomposition (SVD) algorithm. COM position was onset-centered, as described above, but trajectories were not centered to the average trajectory. Note that we also tried the same PCA analysis on non-interpolated data using the alternating least squares algorithm: conclusions were not affected. Nonlinear dimensionality reduction using the *t*-distributed Stochastic Neighbor Embedding (t-SNE) was performed on the same matrix of interpolated and truncated PF trajectories as for the PCA SVD analysis.

For the diffusion analysis (Fig 7B-C), PFs defined on less than 30 laps were excluded. All interpolated PF trajectories (interpolation ensuring that the sample size is constant from lap to lap) were onset-aligned and truncated at 30 laps. The Mean Squared Displacement (MSD) on a given lap was defined as the square of the onset-centered COM position averaged across all PFs. For a random walk in a 1D environment such as our linear track, *MSD* = 2 × *D* × *Lap* #, where D is the diffusion coefficient (Einstein, 1905). In other words, COM shifts can be described by a random walk when D is constant, i.e. when the MSD is a linear function of post-onset lap. D was assessed by linear regression of the MSD as a function of lap number, using the Matlab *regress* function (excluding the first 3 laps, where we observed large nonlinear changes in MSD). Alternatively, to avoid assumptions about when the relationship becomes linear, we assessed the instantaneous diffusion coefficient D_n_ of lap n (equation 3) and fitted it with a decaying exponential to estimate D as the asymptote value of D_n_ (equation 4):

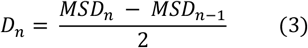

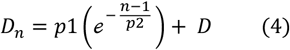

Parameters p1, p2 and D were optimized using the Matlab *fit* function with the nonlinear least square method and Trust-Region algorithm. The parameter search starting point was [100, 2, 0]. Parameter search was bounded such that 0 ≤ p1 ≤ 1000, 0 ≤ p2 ≤ 100 and 0 ≤ D ≤ 20.

### PF width

Throughout the study, PF width was characterized by the “standard deviation” (sd) of PF activity:

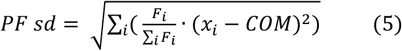

Where i is the spatial bin index on the track, F_i_ is either the normalized fluorescence (for experimental data) or firing rate (for simulated data) in bin i, x_i_ is the position of bin i, and COM is the PF center of mass. PF width was the PF sd of PF activity averaged across all laps. We also computed the lapwise PF sd and assessed the change in width (*PF* ∆ *Width*) as the difference between the first 3 laps and the last 3 laps.

### Place cell model with plastic synapses following an STDP rule

To simulate experiments like in Dong et al. (2021), we considered a virtual mouse running unidirectionally on a 300 cm linear track at constant speed for 30 laps (note that the unidirectional motion with immediate teleportation from end to start makes it equivalent to a circular track, as in Yu et al. (2006)). We designed a simple feedforward place cell model (Fig 2A) that consisted of a leaky-integrate-and-fire (LIF) output neuron receiving weighted synaptic inputs from N spatially modulated input neurons, with one synapse per input neuron. Each input neuron generated spikes stochastically based on a nonhomogeneous Poisson process governed by a single Gaussian place field defined by its COM, peak firing rate (Peak FR_in_) and width (PF_in_ sd). The COM of input PFs regularly tiled the length of the track, and the initial connectivity vector followed a Gaussian defined by its standard deviation (connectivity sd) and a maximum synaptic weight 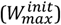 for the input neuron with COM in the middle of the track. In this model, an input spike from neuron j results in an excitatory post-synaptic current (EPSC) with maximum amplitude at the time of the spike defined by the current synaptic weight, w_j_(t), of synapse j. EPSCs then exponentially decay with time constant *τ*_*EPSC*_. The input current I(t) to the LIF output neuron was computed based on the following ordinary differential equation (ODE):

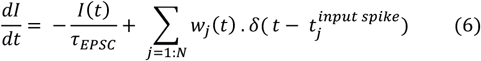

Where 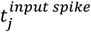 is the time of an input spike at synapse j and δ is the Dirac delta function (1 at 0 and 0 otherwise).

The membrane potential V_m_ of the LIF output neuron was governed by the following ODE (Dayan & Abbott, 2005):

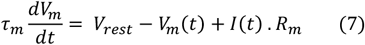

Where *τ*_*m*_ is the membrane time constant, V_rest_ is the resting membrane potential and R_m_ is the membrane resistance. Each time V_m_ reaches the spiking threshold V_thresh_, an output spike is fired and V_m_ is reset to V_reset_.

In Fig S3 and S4 we added spike rate adaptation to the LIF equation using an additional SRA variable that exponentially relaxes to 0 after an increment of potassium leak current at each new output spike, as described in Dayan and Abbott (2005):

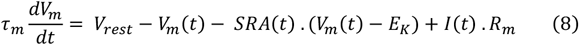

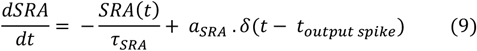

Where E_K_ = -70 mV is the equilibrium potential of potassium, *a*_*SRA*_= 0.06 is the increment value for the SRA variable, and *τ*_*SRA*_ = 100 ms is the time constant controlling the decay of the SRA variable. Parameter values were as in the example provided in Fig 5.6 in Dayan and Abbott (2005).

The synaptic weight of each synapse evolved independently following an antisymmetric pair-based spike-timing-dependent plasticity rule (Fig 2B) where the weight of synapse j potentiates or depresses depending on the delay between a pre-synaptic spike and a post-synaptic spike as follows:

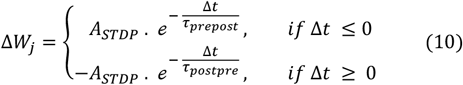

Where ∆*W* is the change in synaptic weight, *A*_*STDP*_ is the maximum amplitude that ∆*W* can take, *τ* _*prepost*_and *τ*_*postpre*_ are the time constants of the exponential decay, and 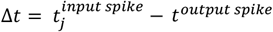 (referred to as the pre-post delay in the rest of the study). Synaptic weights were updated additively using local eligibility variables for each input and output neurons (Morrison et al., 2008; Song et al., 2000; X. Yu et al., 2006). For a given synapse j, the pre-before-post variable P_prepost_ (corresponding to a negative pre-post delay and thus to the potentiating portion of the STDP rule) is triggered on input spike times and decays with time constant *τ*_*STDP*_, whereas the post-before-pre variable P_postpre_ (corresponding to a positive pre-post delay and thus to the depressing portion of the STDP rule) is triggered on output spike times, decaying with the same time constant since the rule is antisymmetric. Weights were updated at each input and output spike times, evaluating P_prepost_ on output spike times and P_postpre_ on input spike times (see Fig 2C). Weight dynamics thus evolved as follows:

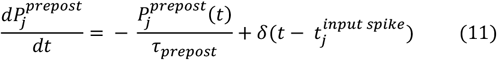

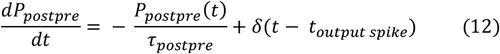

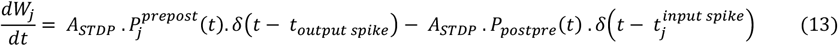

Weights were updated instantaneously unless otherwise stated (as shown in Fig 2C). Because this is not realistic and that a previous model resulting in PF backward shifting implemented a delay (at the end of each lap) in the update (Mehta et al., 2000), we added some dynamics to the weight update in some simulations (Fig S3-4). We designed a phenomenological model where the target weight W_target_ is set by equation 13, and the true weight W exponentially adjusts to that target with a time constant *τ*_*update*_ of 5 seconds (see Parameterization) based on equation 14:

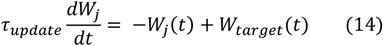

Throughout, ODEs were solved using Euler’s forward method, with a time step of 1ms. Initial conditions: V(0) = V_rest_, all other variables started at 0. Synapses were saturating unless otherwise stated: weights were hard-bounded by EPSC_min_ (0 pA) and EPSC_max_ (same value as 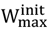). The baseline parameters, corresponding to model 1 in Fig 2C, are shown in Table 1. Alternative parameters are directly mentioned in the figures and legends.

**Table 1.** Baseline parameters.

| Parameter | Value | Parameter | Value | Parameter | Value |
| --- | --- | --- | --- | --- | --- |
| **<u>Experiment</u>** |  | **<u>Output</u>** |  | **<u>Synaptic plasticity</u>** |  |
| Animal Speed | 15 cm/s | $R_m$ | 100 MOhms | Saturating synapses | yes |
| Track length | 300 cm | $\tau_m$ | 20 ms | $\text{EPSC}_{\text{min}}$ | 0 pA |
| Number of laps | 30 | $V_{\text{rest}}$ | -70 mV | $\text{EPSC}_{\text{max}}$ | 85 pA |
| **<u>Inputs</u>** | | $V_{\text{thresh}}$ | -54 mV | $\tau_{\text{prepost}}$ | 20 ms |
| N | 100 input neurons | $V_{\text{reset}}$ | -60 mV | $\tau_{\text{postpre}}$ | 20 ms |
| Peak $\text{FR}_{\text{in}}$ | 10 Hz | SRA | no | $A_{\text{STDP}}$ | 0.5% of $\text{EPSC}_{\text{max}}$ |
| $\text{PF}_{\text{in}}$ sd | 18 cm | | | Weight update | instantaneous |
| Connectivity sd | 10 input neurons |  |  |  |  |
| $W_{\text{max}}^{\text{init}}$ | 85 pA | | | | |
| $\tau_{\text{EPSC}}$ | 10 ms | | | | |

**Table 2.** Comparison of our baseline model with seminal studies. PF width was reported as half-max in the original studies, which we converted to PF sd for comparison (sd = half-max width / 2.355).

|  | **Mehta et al. 2000** | **Yu et al. 2006** | **Model 1 (Fig 2C)** |
| --- | --- | --- | --- |
| Nb of simulations | not reported | not reported | 100 |
| Nb of laps | 20 | 20 | 30 |
| Track Length (cm) | 200 | 200 | 300 |
| Speed (cm/s) | 50 | not reported | 15 |
| **Inputs** |  |  |  |
| N (nb of inputs) | 100 | 1000 | 100 |
| Neuron Type | Rate<br>(gaussian current) | Poisson spiking<br>from gaussian rate | Poisson spiking<br>from gaussian rate |
| PF<sub>in</sub> sd (cm) | 12.7 cm | 3, 12.7 and 30 cm | 18 cm |
| Peak FR<sub>in</sub> | \ | not reported | 10 Hz |
| Initial Connectivity | Gaussian -<br>unreported SD | Gaussian or Skewed -<br>unreported SD | Gaussian: SD = 10 i.n. |
| Synaptic Efficacy<br>(unit) | EPSC (Amps) | unitless G | EPSC (Amps) |
| Max input | 1 nA | not reported | 85 pA |
| **Output** |  |  |  |
| Neuron Type | Rate | Spiking<br>(LIF, parameters<br>unclear) | Spiking<br>(LIF, Song et al. 2000<br>parameters) |
| Spike-Rate<br>Adaptation | Yes<br>(Wang 1998<br>method) | No and Yes<br>(Dayan & Abbott<br>method) | No |
| Inhibition | Oscillatory (8Hz) | Divisive | No |
| Peak FR<sub>out</sub> | high<br>(~50Hz on lap 1) | high<br>(~ 50 to 100Hz) | realistic<br>(~10 Hz) |
| **Plasticity** | STDP | STDP | STDP |
| Implementation | lap integration | local eligibility<br>(Song et al. 2000) | local eligibility<br>(Song et al. 2000) |
| A<sub>STDP</sub><br>(Max Potentiation) | 0.6% of EPSC<sub>max</sub><br>= 0.6 pA | 0.5% of G<sub>max</sub> | 0.5% of EPSC<sub>max</sub><br>= 0.425 pA |
| Minimum<br>Depression | -0.9A<sub>STDP</sub> = -0.54 pA | -0.525% of G<sub>max</sub> | -0.5% of EPSC<sub>max</sub><br>= -0.425 pA |
| Time Constant<br>(ms) | 10 | 20 | 20 |
| Saturating<br>synapses<br>(Upper Bound) | No | Yes<br>(not reported) | Yes<br>(hard bound : 85 pA) |
| Weight Update | end of lap | instantaneous | instantaneous |

### Parametrization

Virtual animal speeds (15 or 25 cm/s) generally corresponded to realistic individual average speeds in mice experiments (Dong et al., 2021; Milstein et al., 2021). 50 cm/s speed was also tested to compare to Mehta et al. (2000), which modeled rats, but is unrealistically high for mice.

The parameters for the output LIF neuron were taken from Song et al. (2000). They correspond to generic cortical pyramidal cell parameters and are within the range of observed values for CA1 pyramidal neurons (Kowalski et al., 2016; Tripathy et al., 2014) (see https://neuroelectro.org/neuron/85/).

Input parameters were chosen to obtain CA1-like output PFs, with realistic width and firing rates (Fig S1). In mice, the median output Peak FR_out_ in dorsal CA1 is ∼10Hz (Mou et al., 2018) and the median PF sd is 13.5 cm in the Dong et al (2021) dataset. Realistic ranges are shown in Fig S1. We used inputs with gaussian PFs inspired from CA3 recordings, but they can also be understood as an average of all spatial inputs to a pyramidal cell, including from the entorhinal cortex (Li et al., 2023; Solstad et al., 2006). PF_in_ sd was chosen to be close to the median value that we observed in CA3 (Fig S1). Peak FR_in_ matches reports from rats in CA3 (H. Lee et al., 2015; I. Lee et al., 2004) which are very close to firing rates observed in the CA1 of rats and mice (I. Lee et al., 2004; Mou et al., 2018). *τ*_*EPSC*_ is within the range of observed values in CA pyramidal cells (Kowalski et al., 2016). The number of input neurons (i.e. synapses) was like in Mehta et al. (2000), and the connectivity sd and maximum initial weight 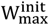 were adjusted to obtain CA1-like PF_out_ sd and Peak FR_out_ as defined above.

We also performed simulations with 1000 input place cells (with connectivity sd at 100 i.n. and 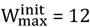), which is a more realistic number of inputs and was used in other models (D’Albis et al., 2015; X. Yu et al., 2006), but results were similar and we thus kept 100 inputs as our baseline for computation speed. Although 100 input place cells is not realistic, note that the distribution of synaptic weights with 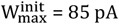 fits well with the amplitude of CA1 EPSPs recorded in vivo or in vitro: EPSPs in vivo are 1.4 mV on average (Kowalski et al., 2016), EPSPs evoked by Schaffer stimulation in slices were ∼2 mV on average (Bittner et al., 2017) which corresponds to 77pA with our LIF parameters (Fig S8), dual patch experiments between CA3 and CA1 pyramidal cells yield EPSCs of similar amplitudes (Dürst et al., 2022) and miniature EPSCs from a single synapse are 15 pA on average (0 to 30 pA range) (Forti et al., 1997).

In Fig 2F and S5, PF_in_ sd, Peak FR_in_ and connectivity sd were varied systematically within a realistic range for CA3 but to cover both realistic and unrealistic PF properties for CA1. We did not vary 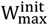, which also controls the output firing rates, because in our model it also conditioned the absolute maximum weight change and we wanted to determine the effect of output rates without changing the amplitude of STDP. STDP parameters (A_STDP_ and time constants) were varied independently of input parameters in Fig S6-7.

STDP parameters were inspired from Song et al. (2000). First, to maintain PFs with realistic Peak FR_in_, synapses were saturating with an upper bound of synaptic weights 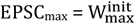 unless otherwise noted (Fig S3-4). Concerning the amplitude of weight changes, although most STDP experiments report them relative to the initial weight of the recorded synapse and thus assume that synaptic modifications depend on the synaptic weight, we considered an additive weight update scheme where A_STDP_ is a constant, like in Song et al (2000) and Yu et al. (2006). We made this choice for several reason: 1) for simplicity and comparison with past models, 2) the weight-dependency of STDP is not clear (Morrison et al., 2007, 2008), especially given that initial weight is just one of many factors potentially influencing long-term synaptic modifications and generally not taken into account by a single STDP rule (Buchanan & Mellor, 2010; Inglebert et al., 2020; Shouval et al., 2010; Wittenberg & Wang, 2006), 4) The additive scheme is a reasonable approximation, especially in the range of EPSCs used in our model (Bi & Poo, 2001; Morrison et al., 2007, 2008), and 5) if synaptic modifications were weight-dependent, an additive scheme like ours would slightly overestimate the effect of STDP for small initial weights, and thus overestimate, not underestimate, the effect of STDP on PF shifting. The baseline value for A_STDP_ was thus set at 0.5% of the maximum synaptic weight, like in Song et al. (2000). However, note that in contrast to Song et al. (2000) and Yu et al. (2006), synaptic weights were defined as EPSCs amplitudes, not unitless conductances. In our model, the baseline absolute maximum weight change is thus 0.425 pA. The amplitude of weight changes due to single pairs of input-output spikes is difficult to assess (Froemke et al., 2006) but the relative and absolute values that we used was in the range of previous estimates: Bi and Poo (2001) estimate A_STDP_ to be ∼1% of the initial EPSC, and for initial EPSCs between 30 and 100 pA their data show a maximum weight change between ∼0.15 and ∼0.5 pA (Morrison et al., 2008). For different STDP protocols and rules, and for a range of initial EPSCs comprising the value of our EPSC_max_, Wittenberg et al. (2006)’s data suggest A_STDP_ to be ∼0.5% of initial EPSCs like we used. To make sure we were not underestimating the effects of STDP, we also explored a range of A_STDP_ values in Fig S6-7: 0.5%, 1%, 2%, 4% or 10% of EPSC_max_, i.e. 0.425, 0.85, 1.7, 3.4 and 8.5 pA (most of these values being outside a realistic range). We did similarly for STDP’s time constants and explored a range of values in Fig S6-7 including the usual estimates (10 or 20 ms) and up to unrealistic values (100 ms).

For models including a synaptic update with dynamic delay, the value of *τ*_*update*_ (5s) was not optimized but grossly corresponds to the dynamics of the early expression phase of long-term plasticity (Gustafsson et al., 1989) and is consistent with the second-timescale of the calcium-dependent enzymatic activation controlling the rapid surface diffusion of AMPA-receptors necessary for the earliest-phase of LTP (Penn et al., 2017; Rodrigues et al., 2021).

Comparison of our baseline model with seminal models of backward PF shifting using STDP can be seen in Table 2.

### BTSP model

The model described above was adapted to have BTSP rather than STDP as the plasticity rule (baseline parameters of table 1 were used unless otherwise stated). BTSP is known to be triggered by a dendritic plateau-potential resulting in a large depolarization with a somatic burst of spikes also called a complex spike (CS)(Bittner et al., 2015, 2017; Cohen et al., 2017; Milstein et al., 2021). Because the mechanisms leading to a CS and triggering BTSP are not well understood, we opted to model BTSP-triggering events simply as a special subset of output spikes, with each regular output spike having a probability p(CS) to be labelled as a CS. The BTSP rule was defined as a pure potentiation rule, as reported in Bittner et al. (2017), with the following kernel (see Fig 3A):

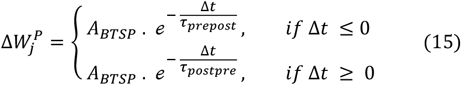

Where 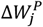 is the potentiation at synapse j due to BTSP, and A_BTSP_ is the maximum potentiation. In order to avoid runaway potentiation and maintain a place field, as observed in Milstein et al. (2021), synaptic weights were not bounded like for the STDP model but obeyed a simple homeostatic rule keeping the total sum of weights constant at each time step. We implemented that homeostatic heterosynaptic plasticity as a weight-dependent synaptic normalization, using a multiplicative scheme (Chistiakova et al., 2015; Kim et al., 2020) such that, for all synapses:

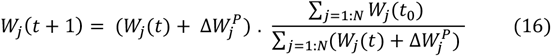

with 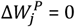 when no potentiation occurred at synapse j.

BTSP-triggered synaptic potentiation was implemented like for STDP (see equations 11-13), using two plasticity variables P_prepost_ and P_postpre_. However, P_postpre_ was not triggered on all output spikes but on CSs, and P_prepost_ was evaluated at the times of CSs only:

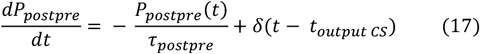

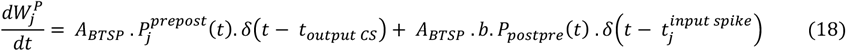

Because there is more temporal summation in the P_prepost_ variable than in P_postpre_, since there are generally more input spikes than output CSs, we added a scaling constant *b* to fit the BTSP kernel on the post-before-pre side (see Validation of the BTSP model and Fig S8).

To avoid making assumptions on the updating dynamics of synaptic weights after BTSP has been triggered (which are not well characterized), we kept the model simple and decided for an instantaneous weight update like for our baseline STDP model. This lack of realism does not impair our conclusions on PF dynamics: most changes due to BTSP are visible on the lap following a BTSP-triggering event (Bittner et al., 2017; Milstein et al., 2021). So, in our simulations, even if the PF activity may be perturbed after a CS on the lap the CS occurred, the PF activity and overall shift will be as expected on the next lap.

Table 3 shows the optimized BTSP parameters used in Fig 3.

**Table 3.** Optimized BTSP parameters.

| Parameter | Value |
| --- | --- |
| **<u>Synaptic Plasticity</u>** |  |
| $\tau_{prepost}$ | 1.31 s |
| $\tau_{postpre}$ | 0.69 s |
| $A_{BTSP}$ | 20 pA |
| $b$ | 1.1 |
| W bounds | no |
| Synaptic Normalization | yes |
| Weight update | instantaneous |

### Validation of the BTSP model

We optimized the BTSP-model parameters to account for the experimental findings from the Magee lab (Bittner et al., 2017; Milstein et al., 2021). BTSP time constants *τ*_*prepost*_ and *τ*_*postpre*_were directly taken from Bittner et al. (2017) (based on the exponential fit of their in vitro dataset). The scaling constant *b* was adjusted by simulating in vitro experiments like in Bittner et al. (2017) so that the maximum potentiation due to BTSP (i.e. without synaptic normalization) would match for both the pre-before-post and post-before-pre part of the kernel and fit the data (Fig S8).

For our homeostatic plasticity rule, we preferred a multiplicative scheme (rather than subtractive) because competition between synaptic resources has been shown to result in such rapid synaptic scaling (Triesch et al., 2018). By design, synaptic normalization operated on the same rapid timescale as BTSP, which is justified on theoretical grounds and has experimental support (Chistiakova et al., 2015).

To optimize A_BTSP_ and to verify that our modeling strategy of combining a BTSP potentiation rule with synaptic normalization yields bidirectional weight changes dependent on the initial weight like observed in vivo in Milstein et al. (2021), we simulated the same kind of experiments and analyzed our resulting dataset in the same way they reported (Fig S9-11). “Milstein-type” experiments (Fig S9-10) consisted in simulating a place cell for 21 laps, with a single CS occurring on lap 11 at a time t_CS_ which was varied systematically to cover the length of the track (there was no relationship between output spikes and the CS in these experiments; t_CS_ was hard-coded). Baseline parameters of our place cell model were used except for the track length (185 cm) and virtual animal speed (25 cm/s) which were as in Milstein et al. (2021). Synaptic weights were updated following the combined BTSP and synaptic normalization rule.

We analyzed subthreshold V_m_ ramps like in Milstein et al. (2021). First, the V_m_ output of the LIF neuron was low-pass filtered (<3Hz) with zero-phase lag (*filtfilt* Matlab function) using a FIR filter with a 2 s Hamming window and wrap-around padding of the V_m_ trace on each lap. For the V_m_ spatial profiles, the low-passed filtered V_m_ traces were binned using 1.85 cm regularly spaced bins and averaged across the 10 laps before or after the CS induction lap. These average traces were smoothed with a Savitzky-Golay filter of order 3 with a window size of 21 spatial bins and wrap-around padding. Temporal profiles of the low-pass filtered Vm (Fig S10C) were binned using the same number of bins as for spatial profiles but not smoothed. The relative amplitude of Vm ramps (used in Fig S10G and S11G) was computed as the difference of the average V_m_ trace with the V_m_ baseline, i.e. Vrest = -70 mV.

Because our goal was to develop a model accurately predicting PF shifting based on BTSP, the optimization objective was to match the high correlation observed by Milstein et al. (2021) between the ramp peak shift and the distance between initial peak and CS, while maintaining a low correlation between pre and post-CS V_m_ (Fig S10-11 and 3C). Our model reproduced key experimental findings, including an apparent weight-dependent bidirectional rule very similar to what Milstein et al. estimated (Fig S10). This rule was computed by linear interpolation of the simulated Vm temporal profiles and the corresponding relative amplitudes of the V_m_ ramps, using the MATLAB *fit* function.

Our approach offers a good fit to the available data on BTSP but is different from past modeling approaches (Cone & Shouval, 2021; Milstein et al., 2021) and has potential shortcomings. First, in our model, only 1 CS is needed to reach a steady-state: adding more induction laps in our Milstein-type simulations does not significantly change the shape of the connectivity vector, which is why we used only 1 induction lap rather than 3 like in the calibration procedure used by Milstein and colleagues for their network model. Whether this one-shot reconfiguration of weights is supported or not by the data is not clear: Milstein and colleagues generally used multiple induction laps, but the number of artificially triggered CSs necessary to induce a new PF was variable (see Fig S1 in Milstein et al. (2021) and figure S7 in (Milstein et al., 2020)) and single spontaneous CSs are sufficient for a new PF to emerge in one-shot (Bittner et al., 2015; Milstein et al., 2021). Note that some of the variability could be due to artificial somatic inductions that may not always trigger calcium plateaus in every dendrite consistently, or not trigger the exact same molecular chain of events than spontaneous dendritic plateaus. More data is needed to clarify how the phenomenology of dendritic plateaus and BTSP co-vary. Similarly, more experiments and analysis are needed to determine whether BTSP-induced depression of the initial PF is slower than emergence of a new one, as predicted by previous models (Cone & Shouval, 2021; Milstein et al., 2021).

The main limitation of our model is that, because synaptic normalization affects all synapses irrespective of the recency of their activity, synaptic potentiation may be underestimated (and depression overestimated) when the CS occurs far from the initial PF. This can result in a relative flattening of the connectivity (Fig S9) and a dilution of the PF activity rather than its translocation, which does not seem to match the Milstein dataset (the maximum increase in Vm was on average larger than the maximum decrease, which was not the case in our simulations). Moreover, connectivity flattening, and thus PF dilution, increase with animal speed (because more inputs are potentiated), making it hard to study the effects of this parameter using our approach. However, despite these limitations, our model fits the data well when CSs occur in-field (Fig S10), which was always the case, by definition, in our in-silico experiments for the study of PF dynamics (Fig 3 and : PF dilution did not occur in these simulations; our model is therefore well-suited to study the effect of BTSP on PF dynamics.

### Statistics, software and hardware

Analyses and simulations were performed using MATLAB (R2021b) on a Dell laptop (Mobile Precision Workstation 3560, i7-1185G7 processor, 16GB RAM, NVIDIA T500 2GB GPU). Statistical details can be found in the legends. In general, we aimed to use estimation statistics as our main line of evidence, emphasizing the effect size and confidence intervals estimates over the significance of p-values (Gardner & Altman, 1986; Ho et al., 2019). Resampling exact tests were used when the sample size was too large for classic hypothesis testing to provide meaningful p-values (i.e. when doing statistics on individual PFs) (White et al., 2014). ANOVAs based on linear mixed-effect models (*fitlme* function) were used for statistics at the level of individual mice, to account for repeated measures (Z. Yu et al., 2022). Bootstrapped estimates and confidence intervals were computed with the *bootci* function, with 1000 bootstrap samples and Bca method. The effect on medians rather than means was evaluated when the sample distribution was not gaussian. Pairwise comparison tests were two-tailed.

## ACKNOWLEDGEMENTS

We thank Aaron Milstein for helpful discussions and Jorge Jaramillo and Lisa Giocomo for their feedback on the manuscript. This work was supported by the NIH (MS: DP2NS111657, AM: F32MH126643), the Whitehall Foundation, the Searle Scholars Program and the Sloan Foundation.

