## Supplemental Figures for "BTSP, not STDP, Drives Shifts in Hippocampal Representations During Familiarization"

**15 Supplementary Figures**

Associated to Fig. 2:

- Figure S1. Place Field width and output firing rates in simulated and recorded place cells
- Figure S2. Model of dynamic input PFs
- Figure S3. More realistic weight update and output firing patterns do not increase PF backward shifting.
- Figure S4. STDP main effect is an unrealistic increase in output rates and PF width.
- Figure S5. Exploration of the parameter space: inputs and animal speed
- Figure S6. Exploration of the parameter space: STDP rules
- Figure S7. Exploration of the parameter space: STDP rules at high input rates and high speed

Associated to Fig. 3:

- Figure S8. Optimization of our BTSP model to match findings from Bittner et al. 2017
- Figure S9. Simulations of experiments as in Milstein et al. 2021: Effect of a single BTSP-triggering event on the pre-existing PF of our place cell model
- Figure S10. A combination of BTSP and synaptic normalization yields an emergent weight-dependent bidirectional plasticity rule and accounts for PF shifts observed in Milstein et al. 2021.
- Figure S11. Characterization of CS-induced PF shifts and emergent weight-dependence of the model used in Figure 3

Associated to Fig. 5:

- Figure S12. Principal components and their distribution for all CA1 and CA3 PFs trajectories combined

Associated to Fig. 6:

- Figure S13. Covariation of the non-linear regression parameters in single PFs
- Figure S14. Cumulative density function estimates of the non-linear regression parameters and the goodness-of-fit for all PFs based on the recorded subfield and environment familiarity.

Associated to Fig. 7:

- Figure S15. COM shifts continue to occur late after onset

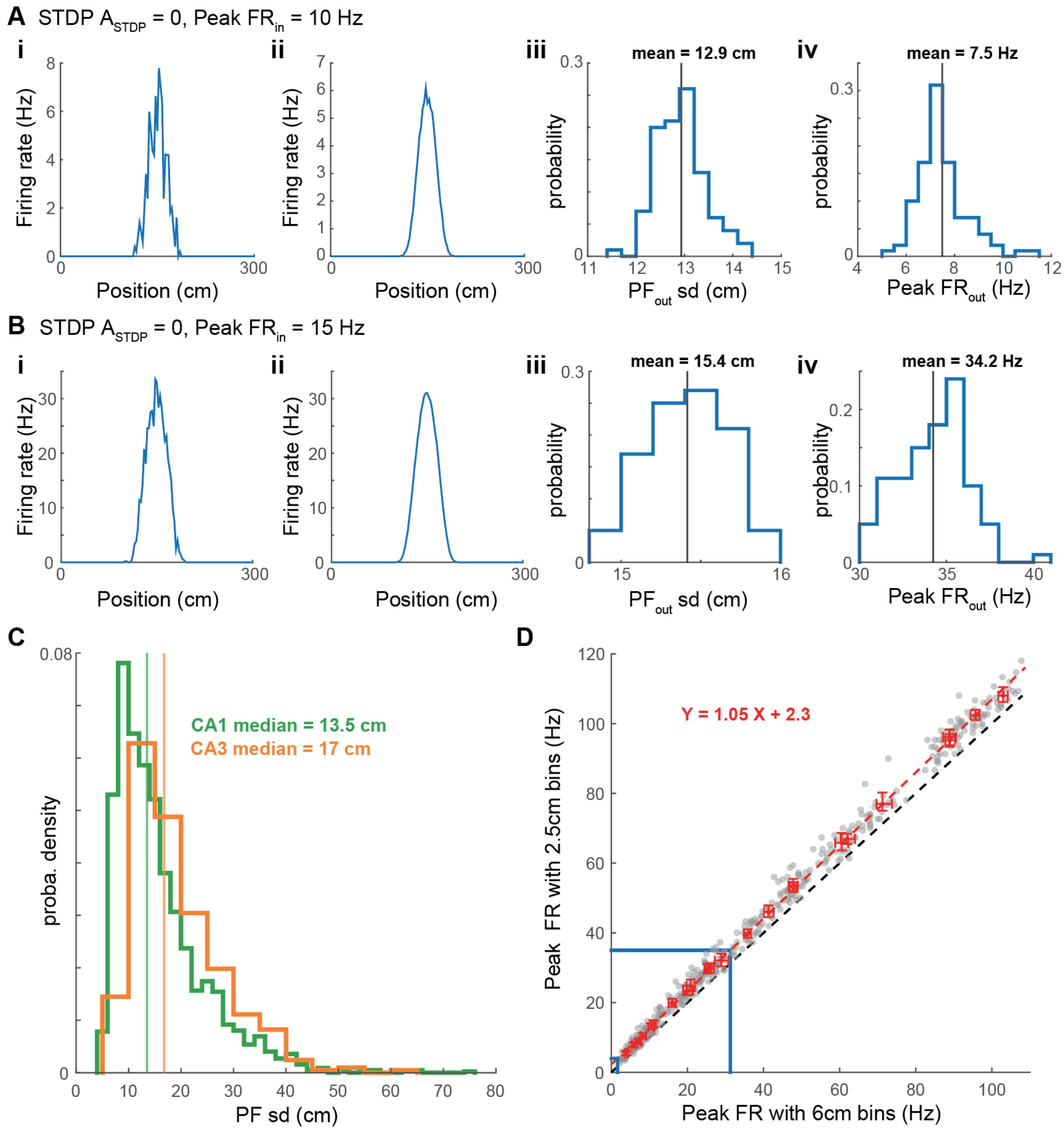


**Figure S1 (associated to Fig. 2). Place Field width and output firing rates in simulated and recorded place cells**

**A-B.** PF models without plasticity, for baseline parameters (A) and higher input rates (B) measured with 2.5 cm spatial bins (like in Mou et al. 2018, but unlike in the rest of the paper which uses 6 cm bins). **i:** Example output PF averaged over 30 laps, **ii**: mean output PF averaged over 100 simulations, **iii**: distribution of output PF width (measured as the standard deviation, sd) for all 100 PFs (30-lap average), **iv**: distribution of peak firing rate of the 100 PFs (30-lap average).

**C.** Distribution of PF width measured in CA1 and CA3 PFs reported in Dong et al. 2021 (same data as in Figure 1). This shows that our model's input PFs (PFin sd = 18 cm unless otherwise stated) and output PFs have realistic width (e.g. see panel A-Biii or Fig. S5B,F).

**D.** Peak firing rates of the 480 simulated PFs in Fig 2Fii, using 6 cm spatial bins (as in Dong et al 2021, Fig 2 and the rest of the study) or 2.5 cm bins as in Mou et al. (2018). Data points are slightly above the identity line (dashed black) showing that using 6 cm bins slightly underestimates firing rates compared to Mou et al.'s methods. Solid blue lines mark the range f Peak Firing rates in data recorded in mice by Mou et al. (2018), and the estimated equivalent when using 6 cm bins based on a linear regression (dashed red). Output PFs from Fig. 2C and S1A-B fall in this range, but not PFs in Fig. 2D.

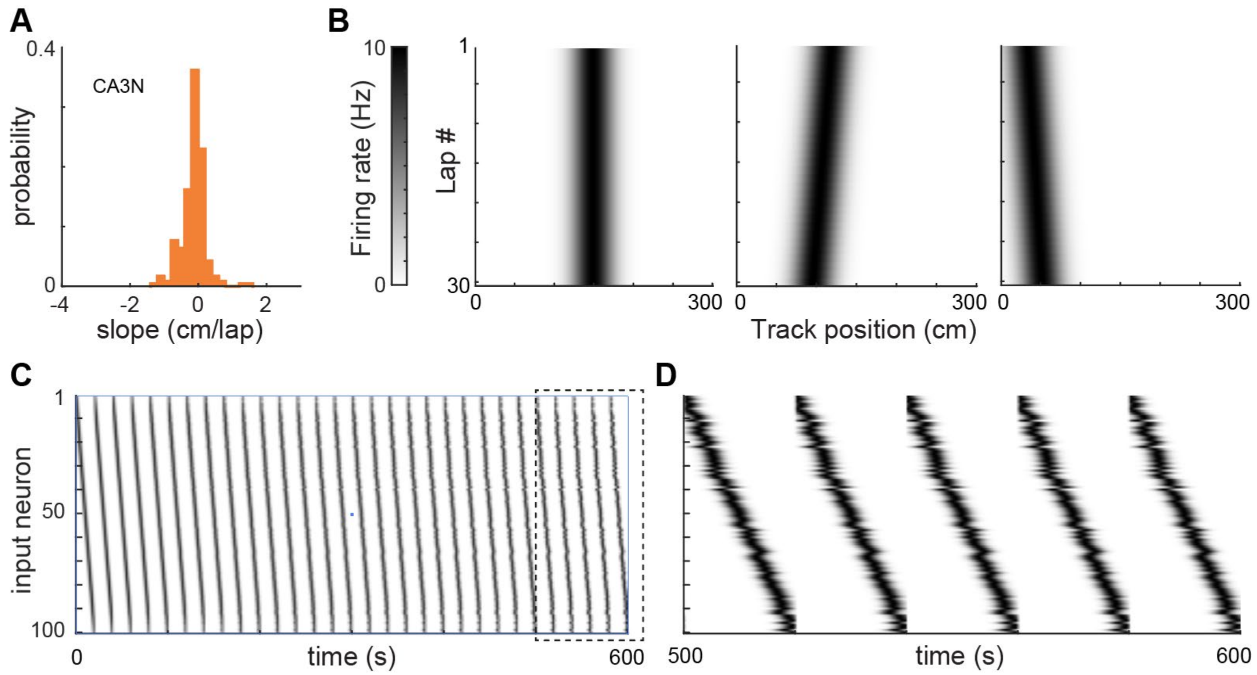


**Figure S2 (associated to Fig. 2). Model of dynamic input PFs**

**A.** Probability distribution of the slope of the linear regression describing the COM shift for each PF recorded in CA3 during exploration of a novel environment (same data as in Fig 1B).

**B.** The lap-wise PFs of inputs to our output LIF neuron were designed to have a shifting slope randomly drawn from the distribution in A (different draw for each simulation). Shown are 3 example input PFs: stable on left, backward shifting in the middle (-0.8 cm/lap) and forward shifting on the right (0.6 cm/lap).

**C.** Input PFs during a full example 30-lap simulation.

**D.** Last 5 laps of the example in C (dashed box).

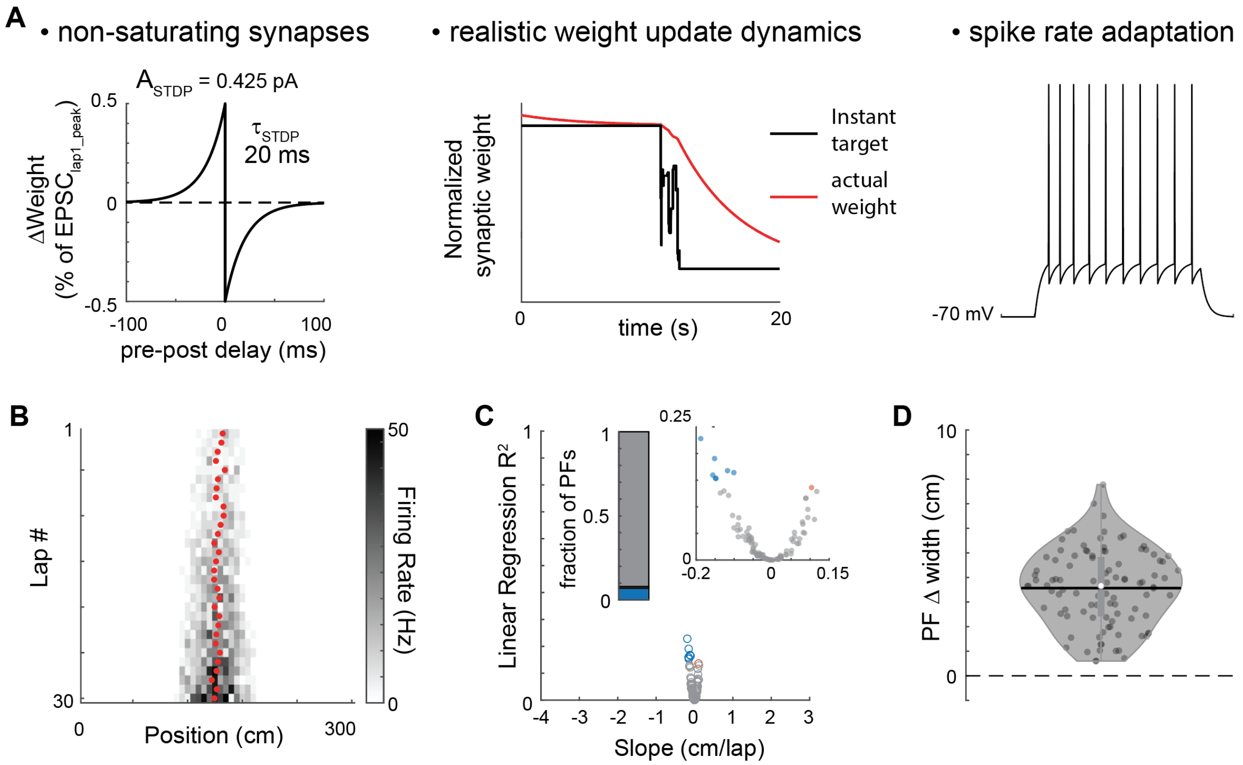


**Figure S3 (associated to Fig. 2). More realistic weight update and output firing patterns do not increase PF backward shifting.**

**A.** The model presented here used baseline parameters except that synaptic weights were allowed to grow without bounds (left: STDP rule), weight update was not instantaneous (middle: example synapse during a single lap) and spike-rate adaptation was added to the LIF output model (right: output neuron response to 500 ms current step of 180 pA). See methods.

**B.** Example simulated PF (not significantly shifting). Lap-wise PF center of mass shown as red dots.

**C-D.** 100 simulations with the same parameters.

**C.** Few PFs are significantly shifting and the shifts are weak. Left inset shows the proportion of PFs backward, forward or not significantly shifting (same color code as Fig 1 and 2). Right inset is the same data zoomed in.

**D.** PF Δ width is computed as the difference of PF sd averaged on the first 3 laps with the PF sd averaged on the last 3 laps. As seen in B, PF width increases for all simulations.

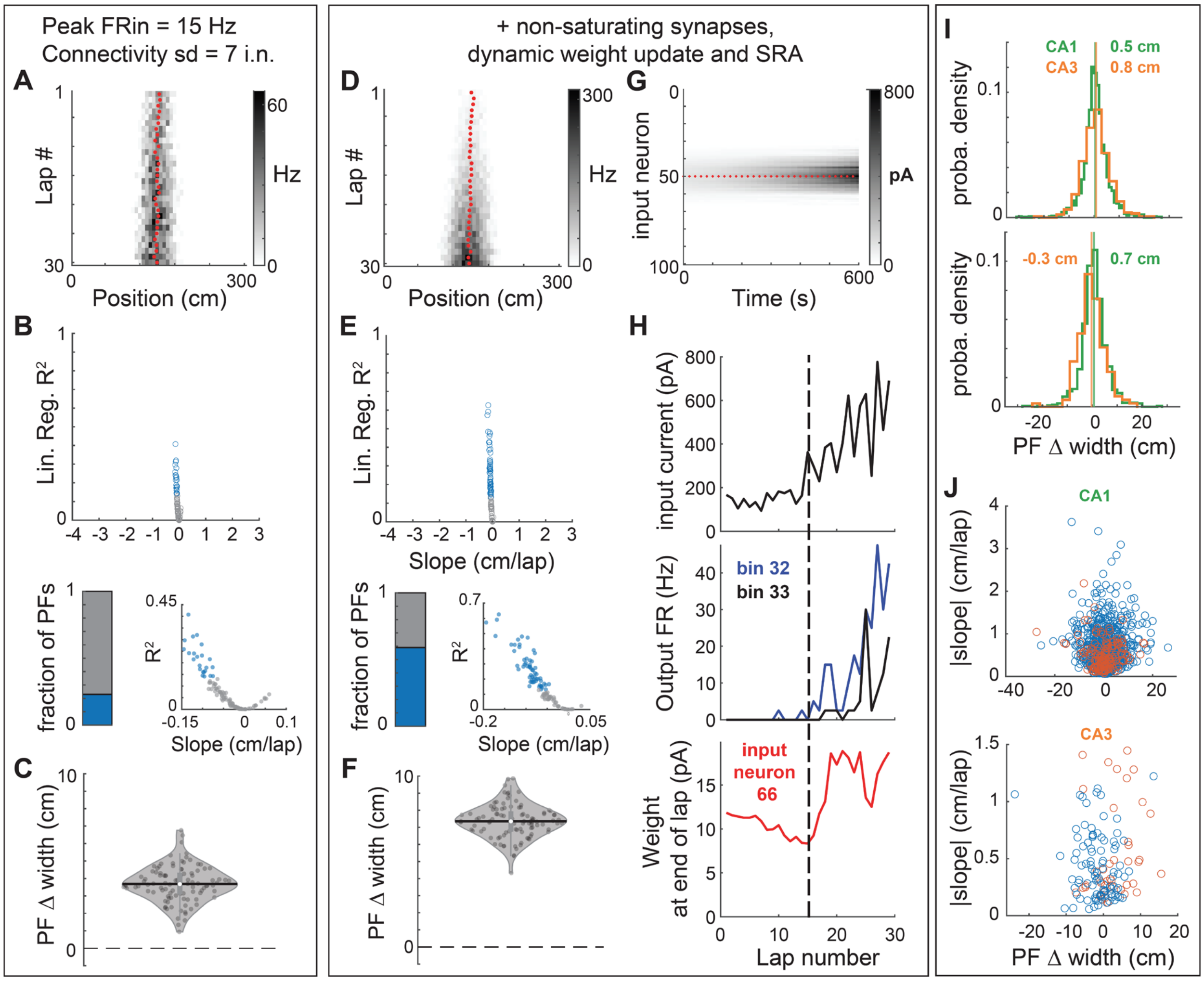


**Figure S4 (associated to Fig. 2). STDP main effect is an unrealistic increase in output rates and PF width.**

**A.** Example simulation with a model with baseline parameters except for a higher input rate (like in Fig 2D) and a narrower initial connectivity vector (this particular set of parameters was also tested in Fig 2F). The PF shows a small backward shift accompanied with an increase in output rates and PF width.

**B-C**. 100 simulated PFs with the same parameters as in A. Only ~20% of PFs show a significant but small backward shift, and no forward shift (**B**). However, all PFs display an increase in PF width (computed as the difference of PF sd averaged on the first 3 laps with the PF sd averaged on the last 3 laps). See Fig S3D for a similar result with different parameters but similar output firing rates.

**D-H.** Same model as in A-C except that, as in Fig S3, synaptic weights were allowed to grow without bounds, weight update was not instantaneous and spike-rate adaptation (SRA) was added to the LIF output model.

**D-F.** Same as A-C. Allowing runaway plasticity leads to an unrealistically large increase in output firing rates (**D**), which yields consistent backward shifting of small amplitude (**E**) and a large increase in PF width (**F**).

**G.** Evolution of the synaptic weight of all 100 input neurons during the 30-lap example simulation in D, illustrating the runaway dynamics (red plus-signs mark the start of a new lap; input 50 correspond to the initial peak weight). Note that the increase in weights is bilateral and happens from inside-out. The COM backward shift is small because weights of inputs > #50 (i.e. with COM in front of the PF center of mass) also increase, albeit slightly slower than inputs < #50.

**H.** The bilateral increase in PF width is due to an inside-out increase in output rates and synaptic weights (same example simulation as in D and G). ***Top***: For each lap, the total input current to the LIF output neuron was averaged over 50 ms preceding the first spike of input neuron #66, which has its PF COM located in bin 33 (the output PF COM being around bin 25). This current corresponds to the sum of inputs < #66 producing the front edge of the output PF. It starts to increase before input 66 weight potentiates (dashed line). ***Middle***: Output firing rate in bin 33 (black) and 32 (blue): note that FR starts to increase in bin 32 (closer to the center) before bin 33, and before input 66 weights start to increase.

***Bottom***: Input 66 synaptic weight depresses first but starts to increase (dashed line) after the output PF has enlarged sufficiently (due to the increased rate in the middle of the PF) to produce output spikes after input 66 spikes. As suggested by past studies, the weights on the forward edge initially tend to depress due to the unidirectional movement and the absence of output firing (thus participating in backward shifting) but STDP makes output rate increase in the center of the PF, which leads to a higher probability of potentiating input-output spike pairs on the edges.
**I-J**. Change in width for PFs recorded in CA1 and CA3 (same data and color codes as in Fig. 1). The distribution is centered around 0, in contrast to models above (**I**: top histogram includes all PFs, bottom only the significantly shifting ones. Vertical lines and associated values are medians). Moreover, PF width changes are not correlated with the shifting speed (**J**: only includes significantly shifting PFs, blue for backward, red for forward).

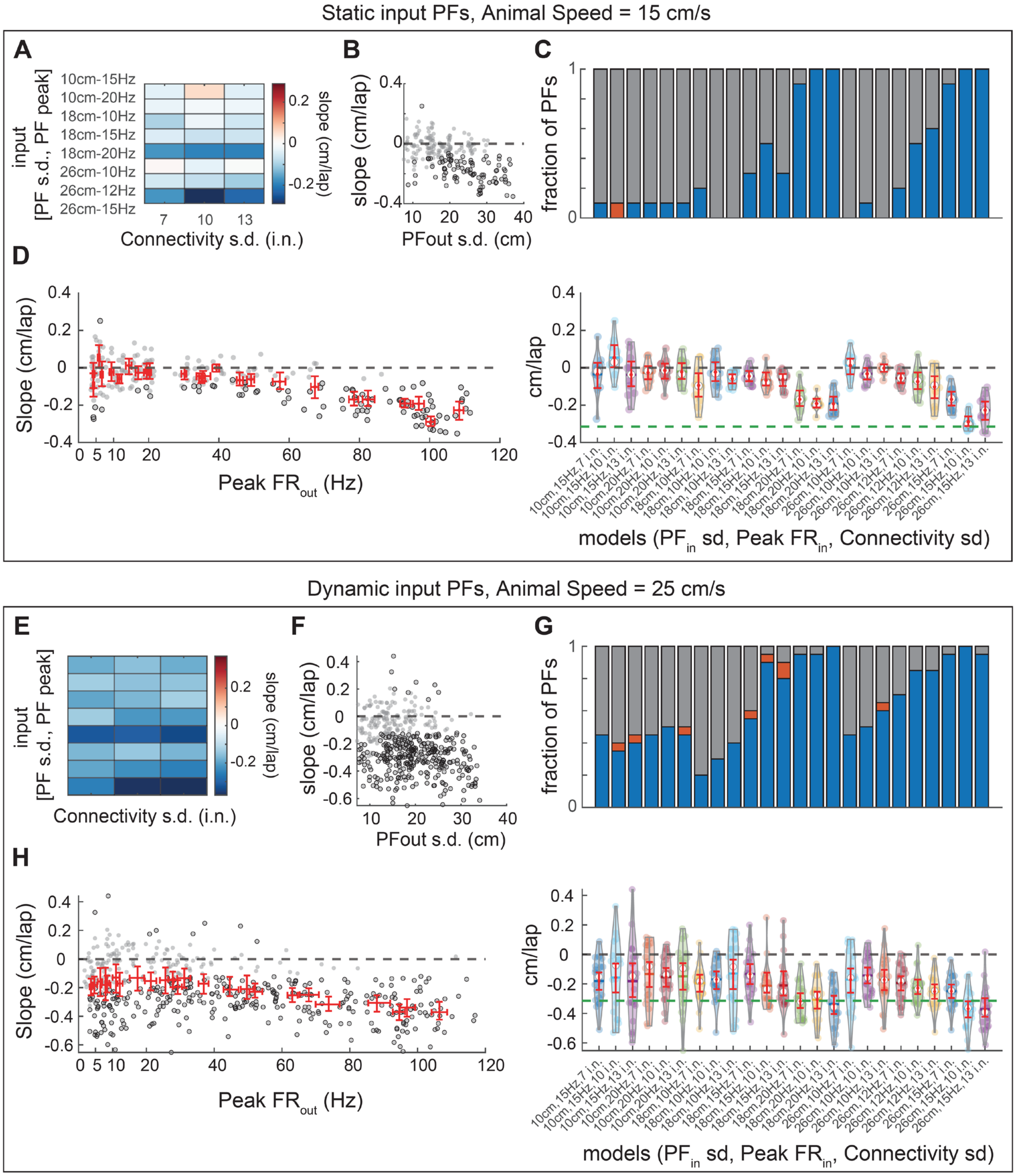


### Figure S5 (associated to Fig. 2). Exploration of the parameter space: inputs and animal speed

**A-D.** Same data as in Fig. 2F.

**A.** Parameters controlling the input rates and width of the initial connectivity vector are systematically varied to explore a wide range (realistic and unrealistic) of output firing rates. For each parameter set, 20 PFs are simulated. The matrix shows for each parameter set the mean COM shifting slope estimated from linear regression (represented as red dots in Fig. 2F and panel D here).

**B.** Shifting slope as a function of output PF width (averaged over the 30 laps). Widths are in a realistic range (see Fig. S1C), with an expected correlation with the slope (significantly shifting PFs artefactually leading to a wider average width).

**C.** Proportion of backward (blue), forward (red) and non-significantly shifting PFs for each set of parameters (Top). The corresponding slope distributions and bootstrapped mean and 95% CI (red) are shown in the bottom panel. The green dashed line marks the mean shifting slope from PFs recorded in CA1 in a novel environment.
**D.** Same plot as right panel in Fig. 2F. It is the same data as in panel C bottom panel, with the same y-axis, but with the peak output firing rate on the x-axis.
**E-H.** Same as A-D but with models including dynamic inputs mimicking PFs recorded in CA3N (see Fig. S2) and with a higher but realistic mouse speed (higher speed promotes backward shifting, because input spikes from farther away can fall in the potentiating time window). Consistent but modest backward shifting (mostly inherited from CA3-like inputs) is seen for all conditions. Some parameter sets yield proportions of backward and forward shifts similar to CA1N but are associated to a mean shift smaller than in real CA1N conditions. Other sets of parameters can yield a mean COM shift similar to CA1N but only because there is a much higher proportion of weakly backward shifting PFs. Moreover, they correspond to unrealistic output firing rates. Overall, no model matched the CA1N dataset well.

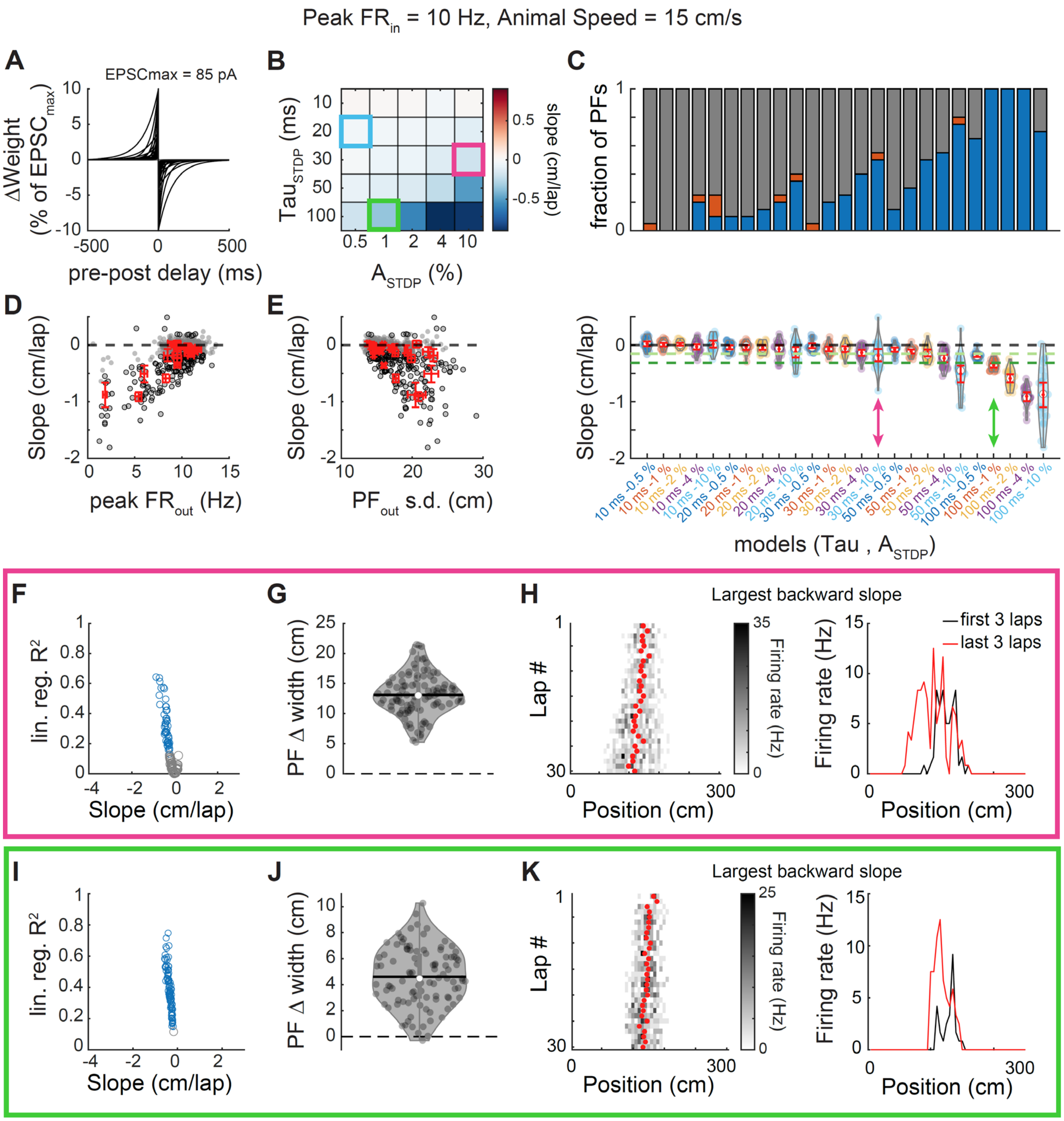


**Figure S6 (associated to Fig. 2). Exploration of the parameter space: STDP rules**

**A.** Different antisymmetric STDP rules were tested, each with a specific time constant Tau (from 10 to 100 ms) and maximum weight change $A_{STDP}$ (0.5%-10% of EPSCmax, i.e. 0.425-8.5 pA). All other parameters were kept as in the baseline model. Note that Tau > 30 ms and $A_{STDP}$ > 2% (i.e. 1.7 pA here) are far from classic STDP rules.

**B.** Mean shift (average across 20 simulations) as a function of the STDP parameters. The light blue square corresponds to our baseline parameter set (model 1 in Fig 2). The pink and green squares highlight 2 sets of parameters that lead to a mean shift similar to CA1F or CA1N, respectively (see panel C bottom).

**C.** *Top*: Proportion of backward (blue), forward (red) and non-significantly shifting PFs for each set of parameters. *Bottom:* The corresponding slope distributions and bootstrapped mean and 95% CI (red). The light and dark green dashed lines mark the mean shifting slope from PFs recorded in CA1F and CA1N, respectively. Pink and green arrows point to the parameter sets highlighted in B, that match the mean shift observed in CA1 (F or N): these correspond to unrealistic $A_{STDP}$ or Tau, respectively, and do not match the range of slopes observed in CA1.

**D-E.** Shifts of all simulated PFs as a function of their average peak output firing rate (**D**) or average width (**E**). Note that large shifts correspond here to a decrease in output firing rates and large PFs (in other words, the PF is diluting away). The low rates and disappearance of the PF are driving the higher variance observed in the most extreme parameter sets.
**F-H**. 100 simulations using the same parameters set highlighted in pink in panels B and C (i.e. that matches the mean shift observed in CA1F). **F.** No forward shift is observed. The range of slope values does not match CA1F. **G.** STDP leads to an unrealistic increase in PF width (see Fig. S4). **H.** Example PF with the largest backward slope in F.
**I-J.** 100 simulations using the same parameters set highlighted in green in panels B and C (i.e. that matches the mean shift observed in CA1N). Here again, the range of slopes, the proportion of backward and forward shifting and the consistent PF enlargement do not match what we observed in CA1N.

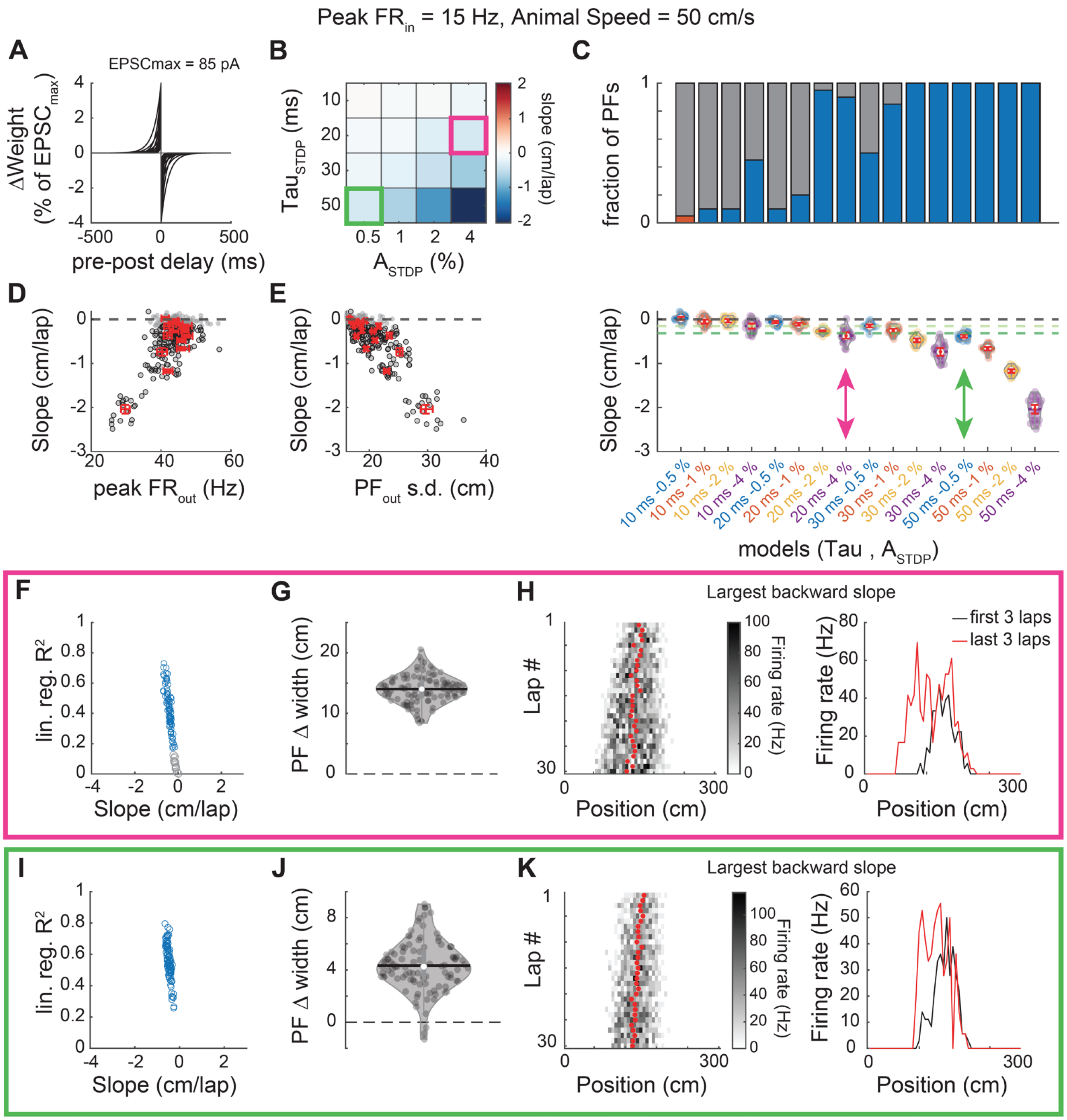


**Figure S7 (associated to Fig. 2). Exploration of the parameter space: STDP rules at high input rates and high speed**

Same as Fig. S6 but with 15 Hz Peak $FR_{in}$ and a virtual animal speed of 50 cm/s (which is unrealistically high for a mouse, but matches the speed used in the model of Mehta et al. (2000), which was inspired by observations in rats. Increasing these parameters promotes the effects of STDP.

**A-E.** Systematic variation of Tau and $A_{STDP}$ (20 simulations per condition). Note that Mehta and Wilson's model used Tau = 10 ms and $A_{STDP}$ = 0.6 pA (which would correspond to ~0.7% of our EPSCmax). Conditions that match the mean shift observed in CA1N are highlighted in pink and green. They exhibit a high proportion of backward shifting PFs unlike what we observed in CA1 and they correspond to unrealistic STDP parameters (high amplitude or high Tau). Backward shifts as large as what we observed in some CA1 PFs (e.g. < - 1 cm/lap, which is not that uncommon, see Fig. 1 and 2E) can only be achieved with unrealistically high STDP parameters.

**F-K.** 100 simulations for each of the parameter sets highlighted in pink (F-H) or green (I-K). The range of slopes, the proportion of backward and forward shifting and the consistent PF enlargement do not match what we observed in CA1.

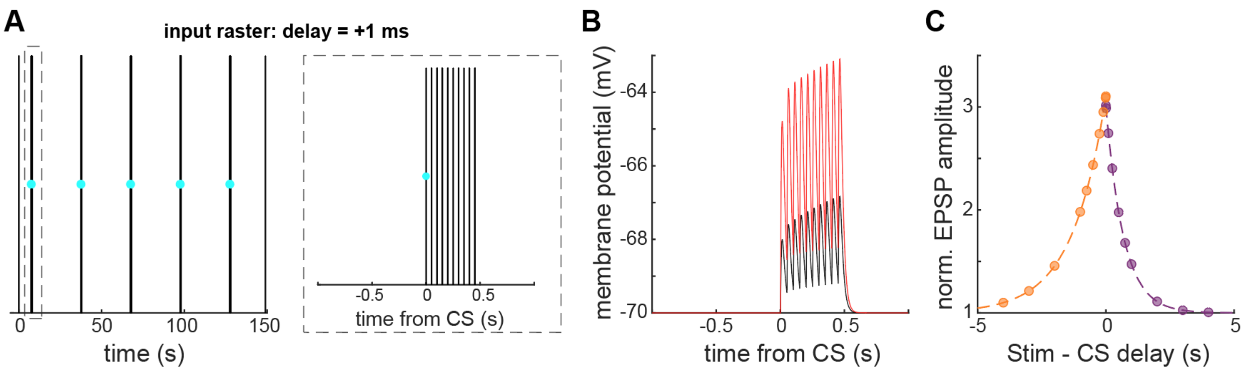


**Figure S8 (associated to Fig 3). Optimization of our BTSP model to match findings from Bittner et al. 2017**

We performed simulations mimicking the experiments done in slices reported in Bittner et al. 2017 (their Figure 3). Our model here consists of the same leaky-integrate-and-fire neuron as elsewhere in our study, but with a single input with initial synaptic weight of 77 pA. This weight yields a 2 mV EPSP when the input fires a spike, which matches the average baseline stimulation-induced EPSP in Bittner et al 2017. BTSP was implemented as a pure potentiation rule (see Figure 3A and S9B) without synaptic normalization (weights were not bounded either), using the time constants estimated by Bittner et al. 2017 (1.31 s for the Pre-before-Post kernel, 0.69 s for the Post-before-Pre kernel).

**A.** The protocol consisted of 5 pairings of a BTSP-triggering complex spike (CS, cyan dot, modeling induction of a plateau in our LIF neuron) with an input spike-train of 10 regularly-spaced spikes at 20 Hz (black vertical lines). The delay between the CS and the beginning of the input train (Stim) was varied from -4 s to 4 s. For a given delay condition, there was an inter-pairing-interval of 30 s. To evaluate changes in EPSP amplitude due to BTSP, single spikes were added 7.5 s before the first CS and 21.5 s after the last CS. Panel A *left* shows an example protocol for the shortest delay (1 ms). The inset on the right shows the first pairing (zoom of the dashed rectangle on the left).

**B-C.** Two parameters were optimized to fit Bittner et al. 2017's results: maximum potentiation = 3.8 pA, scaling factor b = 1.1. The scaling factor value of 1.1 was kept for all BTSP models throughout the study. The 3.8 pA spike-wise maximum potentiation $A_{BTSP}$ fits the in vitro experiments modeled here, but was not kept in models of place cells used elsewhere because the Bittner in vitro experiments were done in conditions with only one input active (i.e. no place field connectivity) and the initial synaptic weights of the stimulated inputs were relatively homogeneous (producing EPSPs ~ 2 mV). The Bittner dataset thus does not allow to determine the effective maximum weight change that would occur in vivo.

**B.** EPSPs during the first (black) and last (red) pairing for the protocol in A, illustrating the potentiating effect of BTSP with the parameters stated above.

**C.** Dashed lines correspond to the BTSP rule determined by Bittner et al. 2017 from their slice experiments. Data points correspond to our simulations (the final EPSP amplitude was normalized to the initial EPSP amplitude, as in Bittner et al. 2017). Compare to Figure 3D in Bittner et al. 2017.

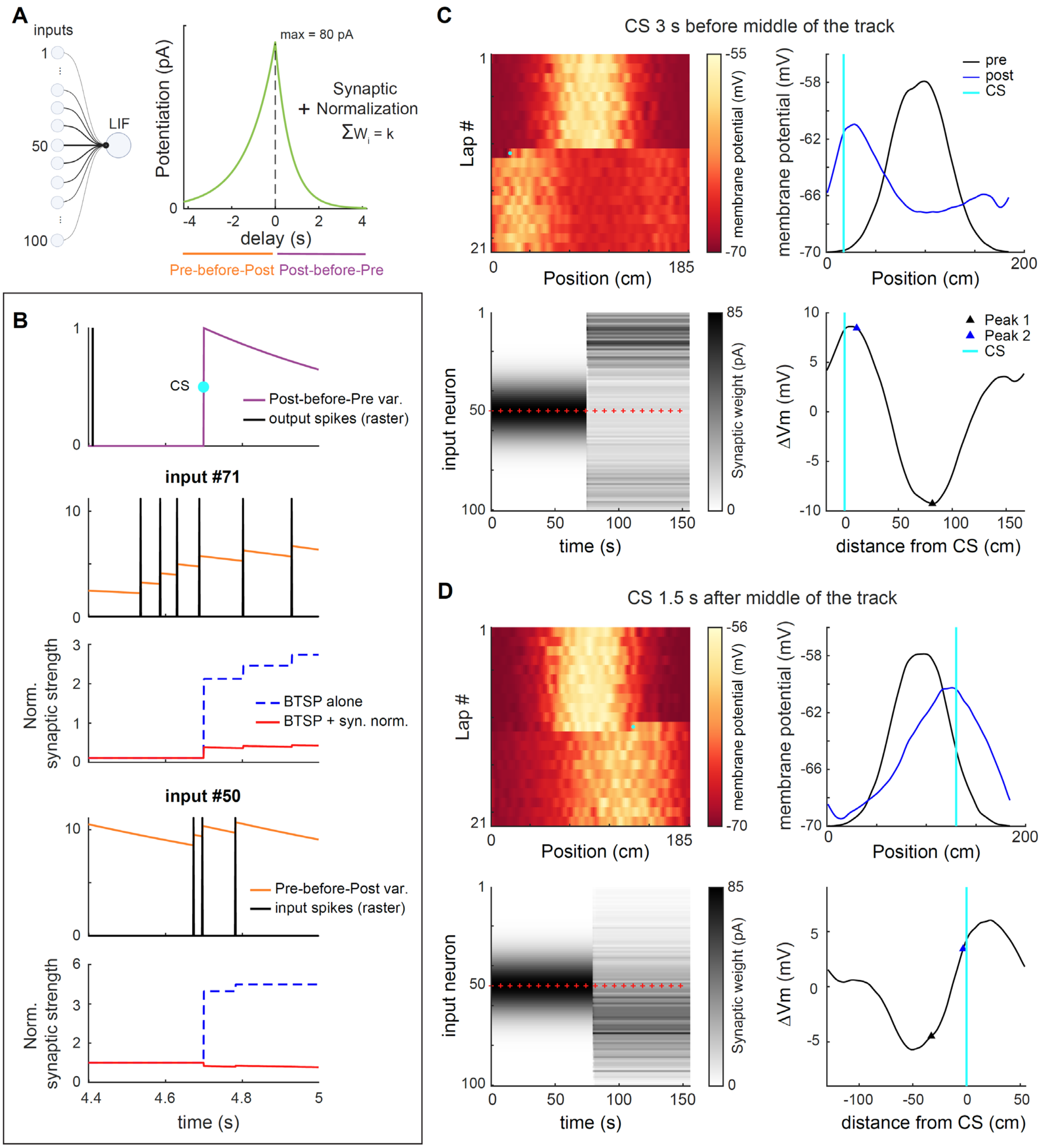


**Figure S9 (associated to Fig. 3). Simulations of experiments as in Milstein et al. 2021: Effect of a single BTSP-triggering event on the pre-existing PF of our place cell model**

**A.** Same place-cell model as in Fig. 2, based on 100 spatially modulated input Poisson processes (with stable PFs) to a single leaky-integrate-and-fire (LIF) neuron. Plasticity was implemented following the potentiation kernel discovered by Bittner et al. 2017 (as in Fig S8) combined with a homeostatic synaptic normalization rule maintaining the sum of weights constant. In the examples shown in this figure, the maximum potentiation of the BTSP rule (i.e. before synaptic normalization) was set at 80 pA.

**B.** Plasticity variables (purple and orange traces) and synaptic weight dynamics (dashed blue and solid red traces) around the time of a complex spike (CS, cyan dot) triggering BTSP, for input 71 (which has a PF close to where the CS occurred) and input 50 (i.e. with PF in the middle of track and maximum initial synaptic weight). Potentiation following the rule in A occurs at the time of CS based on the value of the pre-before-post variable for each synapse (orange), and it also occurs when an input spike coincides with a non-zero value of the post-before-pre variable started by the CS (purple). Normalized synaptic strength is the synaptic weight in pA divided by the maximum synaptic weight of the simulation (85pA – this is not a hard bound here but a consequence of maintaining the sum of weights constant). This example comes from lap 11 in panel D.

**C-D.** Simulations of experiments as in Milstein et al. 2021. Activity in our place cell model is simulated for a virtual animal running 21 laps on 185 cm linear track at a constant speed of 25 cm/s. A CS (cyan dot, BTSP-triggering event) is triggered 3s before the virtual animal reaches the middle of the track (**panel C**) or 1.5s after (**panel D**). *NorthWest*: lapwise membrane potential dynamics (Vm was averaged for each 100 spatial bin). The CS (cyan dot) causes a shift of the spatially modulated Vm. *SouthWest*: Synaptic weight dynamics for all inputs. Red plus-signs mark the start of a new lap. *NorthEast*: Smoothed Vm average for the 10 laps before or after CS. Compare to Figure 1C and S2 in Milstein et al. 2021. *SouthEast*: Difference between the pre-CS and post-CS averages. As in Milstein et al. 2021's Figure S2A and network model optimization procedure, a CS occurring 3s before the middle of the track (i.e. PF center of mass of the input neuron with initial peak synaptic weight) causes a maximum increase in Vm of ~8mV (panel C).

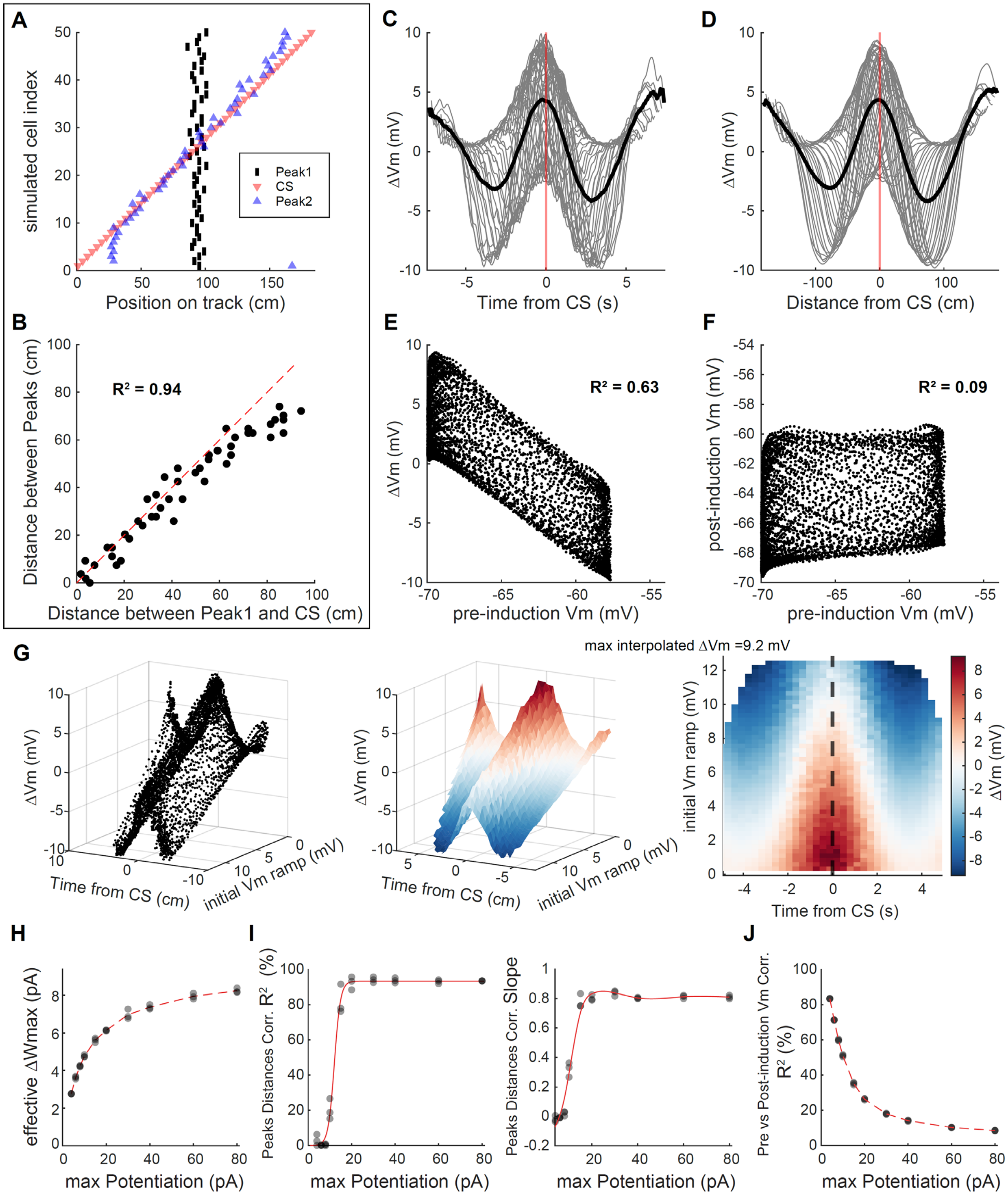


**Figure S10 (associated to Fig 3.). A combination of BTSP and synaptic normalization yields an emergent weight-dependent bidirectional plasticity rule and accounts for PF shifts observed in Milstein et al. 2021.**

**A.** 50 Milstein-type inductions were simulated as in Fig S9, varying the time of CS systematically to span the whole track (185 cm of length, with animal speed of 25 cm/s as in Milstein et al. 2021, and a maximum potentiation of 80 pA before synaptic normalization). Compare to Figure 1D in Milstein et al. 2021 article.

**B.** The distance between Peak1 and Peak2 vs. the distance between the CS and Peak1 are correlated ($p < 0.0001$ computed by Pearson's correlation). Red dashed line is unity, not the regression line. Compare to Figure 1E in Milstein et al. 2021: the PF shifts observed are similar, especially when the CS occurs close enough to Peak1 (which is always the case in our simulations in Figure 3 because, by design, CSs always occur in-field).
**C-D.** Temporal and spatial profiles of CS-induced changes in membrane potential Vm (ΔVm). Grey lines are individual simulations (as in Fig S9C-D South-East) aligned on the CS time or position (red line), the black line is the average of all 50 simulated cells. The spatial profile D is computed from spatially binned and smoothed Vm as in Fig S9C-D North-East. Compare to Figure 2 in Milstein et al. 2021 (note that the x-axes cover a wider range in our case, and that ΔVm beyond 5s away from the CS are not well sampled in the Milstein dataset).
**E.** Changes in Vm are negatively correlated to initial Vm (5000 points, from all 50 simulations in panel A, using binned and smoothed Vm values as in panel D). Compare to Figure 3G in Milstein et al. 2021.
**F.** When the maximum potentiation parameter is high enough (here 80 pA), the correlation between pre and post-CS Vm is poor (low $R^2$). 5000 points from all 50 simulations using binned and smoothed Vm values. Compare to Figure 3H in Milstein et al. 2021.
**G.** Changes in Vm ramp (ΔVm) as a function of both time and initial Vm ramp from baseline (a proxy for initial synaptic weight in Milstein et al. 2021. Indeed, in our model, initial Vm is simply a noisy reflection of initial synaptic weights). *Left*: Same data as in C. *Middle*: Linear interpolation of the data on the left. *Right*: Heatmap of the interpolation, restricted to the range of 'time from CS' in Figure 3I of Milstein et al. 2021. The white band (ΔVm = 0) corresponds to equilibrium, i.e. the target synaptic weight when BTSP is triggered. This analysis shows that our model of BTSP as pure potentiation combined with an additional homeostatic synaptic normalization rule results in an apparent weight-dependent bidirectional plasticity rule like what was suggested in Milstein et al. 2021's Figure 5C.
**H-J.** Varying the maximum potentiation parameter of the BTSP rule affects the fit of the model to Milstein et al. 2021's experimental findings. For each value of the maximum potentiation parameter, we simulated 3 experiments (grey dots) like in panel A (each experiment including 25 simulated inductions, with CSs positions spanning the length of the track).
**H.** The effective maximum weight change (Effective ΔWmax) of the combined BTSP + synaptic normalization model is a non-linearly increasing function of the maximum potentiation parameter of the BTSP rule. A potentiation parameter of 80 pA as chosen in panels A-G corresponds to an effective ΔWmax of ~8 pA, similar to the effective ΔWmax for the parameters that we used in our main in silico experiments in Figure 3. Effective ΔWmax was computed as the maximum weight change in all simulations for a given set of parameters divided by the value of the Pre-before-Post plasticity variable of the given input cell at the time of maximum weight change. Effective ΔWmax is thus an estimate of the maximum weight change for a single input spike, excluding temporal summation, and is comparable to the maximum potentiation parameter of a bidirectional plasticity rule (such as the STDP rule in Figure 2, or Milstein et al. 2021's Figure 5C). However, note that because our model is not designed as a bidirectional plasticity rule but includes a heterosynaptic homeostatic rule, there is no true ΔWmax and the estimate will vary from simulation to simulation. This is because, in our model, weight changes are not only dependent on initial weight but also on the instantaneous sum of weights, and thus on the stochastic activity from all input cells.
**I.** Effect of the maximum potentiation parameter on CS-induced PF shifts characterized as in panel B. Both the strength of the correlation between the CS position and the PF shift ($R^2$, *left*) and the slope of the regression (*right*) reach a plateau at ~20pA, where PF shifts qualitatively fit the Milstein dataset.
**J.** Effect of the maximum potentiation parameter on the strength of the correlation between the pre and post-CS membrane potential, computed as in panel F. This correlation becomes low like in Milstein et al. 2021 for high values of the parameter, such as the 80 pA chosen for panels A-G.

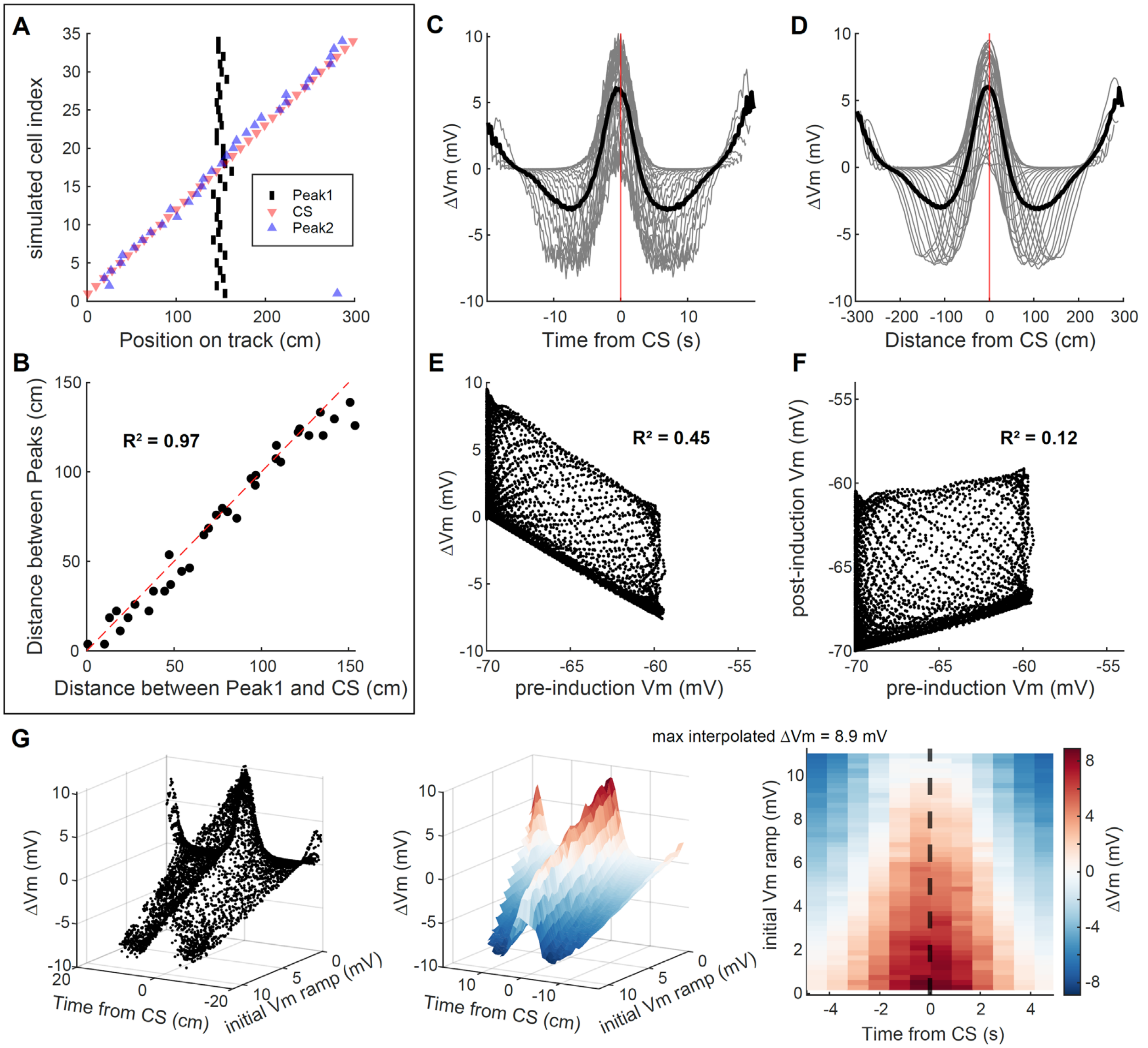


**Figure S11. Characterization of CS-induced PF shifts and emergent weight-dependence of the model used in Figure 3**

**A-G.** Same as in Fig. S10 but for the parameters used in our main in silico experiments shown in Figure 3D-J, i.e. with parameters matching Dong et al. 2021's experimental conditions (track length = 300 cm, animal speed = 15 cm/s) and a maximum potentiation of the BTSP rule = 20 pA (optimized to match Milstein's experimental findings, see Fig. 3C). The analysis was based on the simulation of 35 Milstein-type inductions for which the CS location was varied systematically to span the whole track.

**A-B.** PF shifts are well correlated with the CS location relative to the initial place field, close to the identity line. Even though we don't have a ground truth for these parameters, we took this as the main indicator that our model is consistent with Milstein et al. 2021 experimental findings and can be applied to study the effects of BTSP on PF shifting dynamics.

**C-D.** Temporal and spatial profiles of Vm changes, centered on the CS. Note that the x-axes are wider than in Fig. S10 because the track is longer and the animal speed smaller.

**E.** The correlation between the Vm changes and pre-induction Vm (a proxy for initial synaptic weights) is as strong as in Milstein et al. 2021.

**F.** The correlation strength between pre and post-induction Vm is low, as in Milstein et al. 2021.

**G.** The combination of BTSP and synaptic normalization, with the parameters used in Figure 3, results in an apparent weight-dependent bidirectional plasticity rule akin to the one proposed in Milstein et al. 2021. Note that it is slightly different from what we showed in Fig S10G as it is defined over a wider range of times relative to CS.

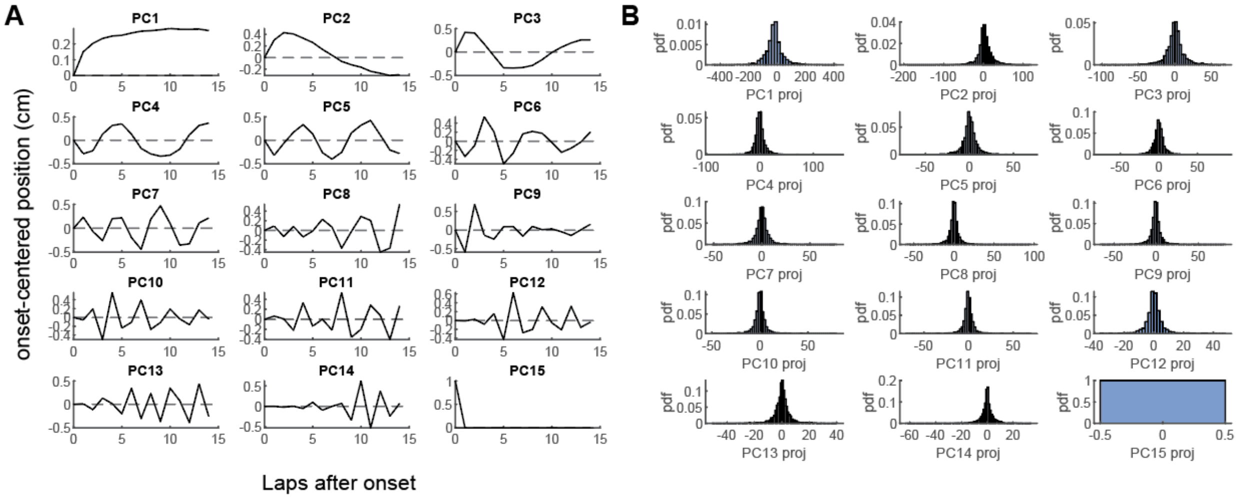

**Figure S12. Principal components and their distribution for all CA1 and CA3 PFs trajectories combined**

**A.** Principal components reveal non-linearities in PF trajectories. See Fig 5B for the explanatory power of each PC.

**B.** PC scores all follow unimodal distributions, revealing no clusters.

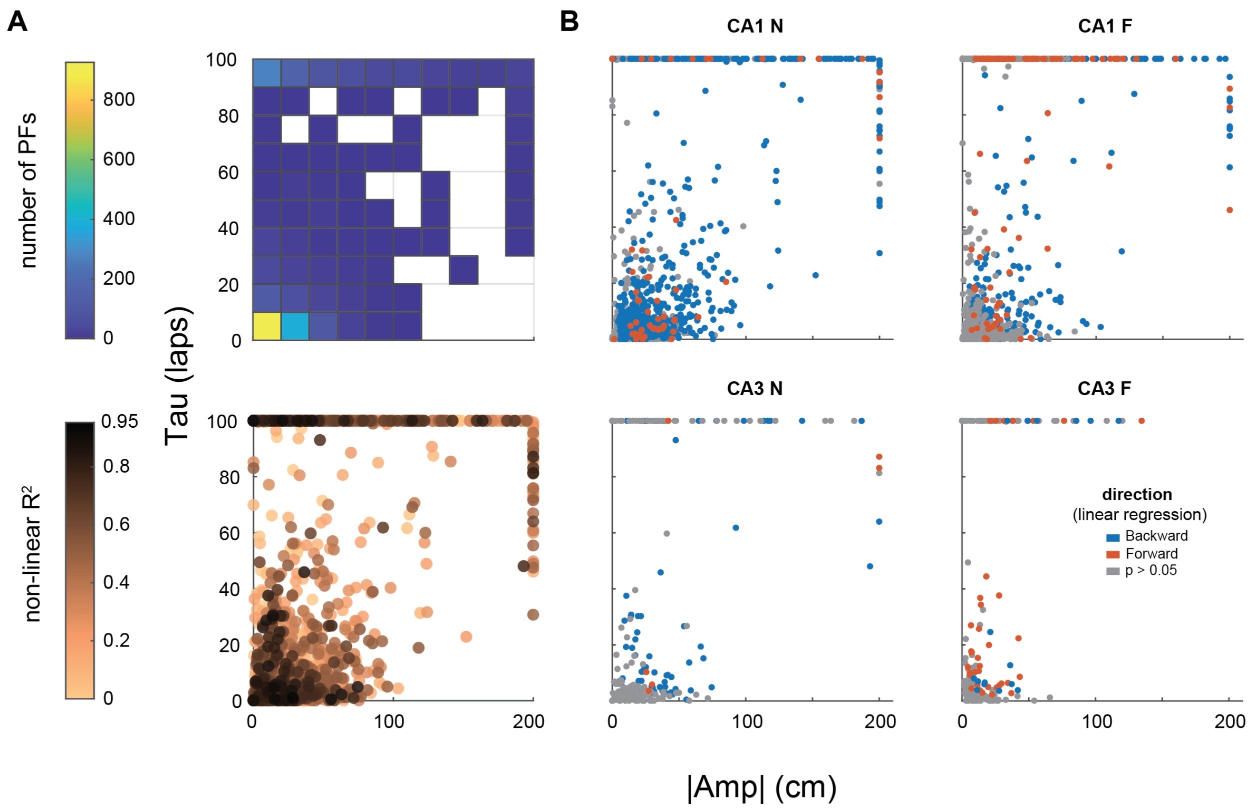


**Fig S13. Covariation of the non-linear regression parameters in single PFs**

Absolute Amp quantifies the amplitude of the COM shift in respect to the initial position. Large tau values correspond to flat trajectories, i.e. linear shifting (when associated to a large Amp) or stable PFs (when associated to an Amp ~ 0).

**A.** *Top*: histogram of the number of PFs in each tile of the parameter space. *Bottom*: corresponding scatter plot, each dot corresponding to a single PF. There is not a clear correlation between Amp and Tau but most PFs are concentrated near the origin, with small Taus (<10 laps) and Amps < 40 cm. There is a second cluster with large Tau values corresponding to stable PFs and to PFs with linear-like dynamics. The nonlinear regression produces good fits ($R^2$) in both clusters. Note that the second group of stable and linear PFs appears as a cluster on the edges because of the fitting procedure that used bounds for |Amp| and Tau at 200 cm and 100 laps respectively.

**B.** Scatter plots of the non-linear regression parameters for each PFs, grouped by recorded subfield (CA1 or CA3) and environment familiarity (N or F).

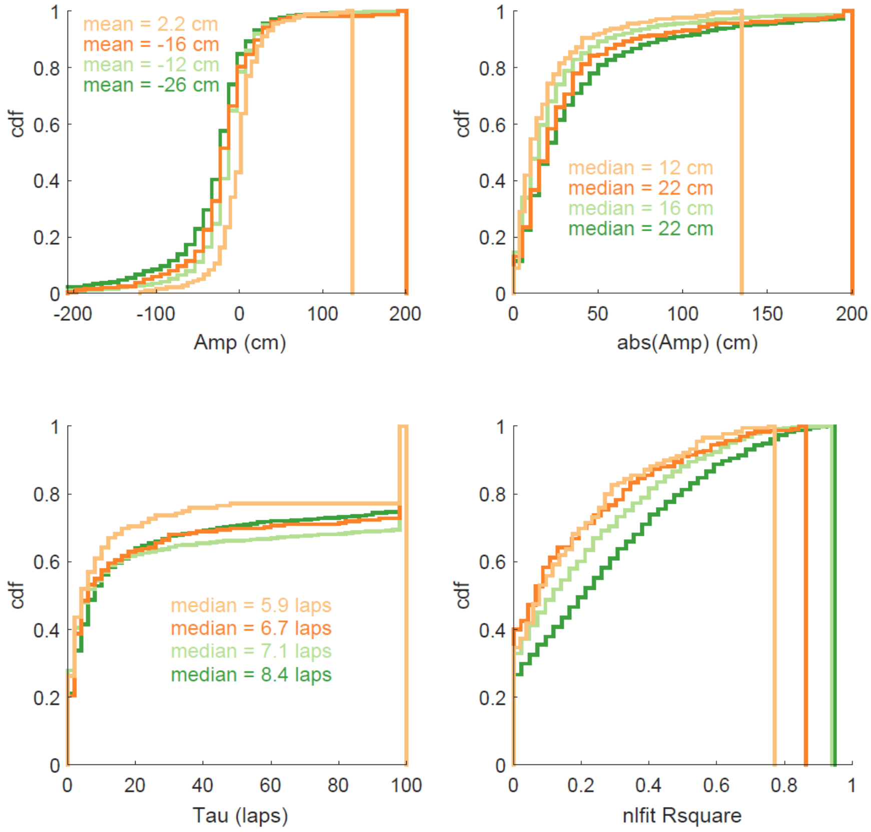

mean = 2.2 cm
mean = -16 cm
mean = -12 cm
mean = -26 cm
cdf
Amp (cm)
median = 12 cm
median = 22 cm
median = 16 cm
median = 22 cm
abs(Amp) (cm)
median = 5.9 laps
median = 6.7 laps
median = 7.1 laps
median = 8.4 laps
Tau (laps)
nlfit Rsquare

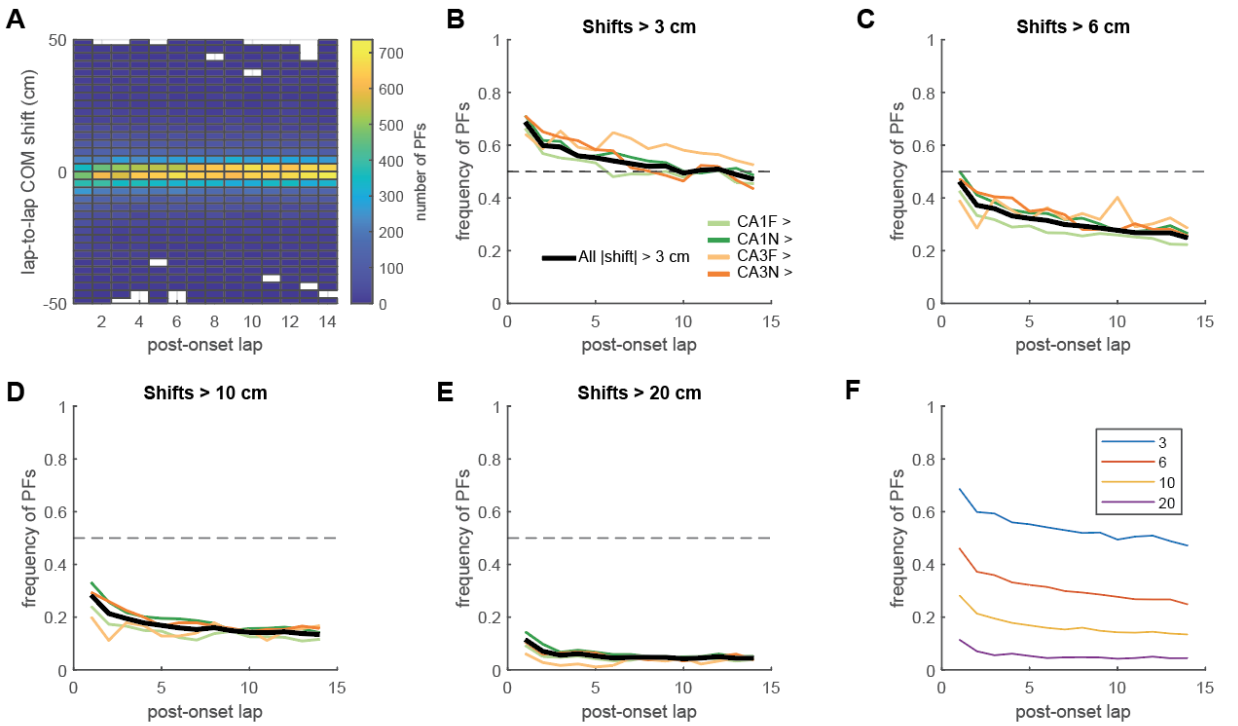


**Figure S15. COM shifts continue to occur late after onset**

Unsupervised analysis designed to detect whether COM shifts continue to occur long after PF onset. **A.** Distribution of $lap_{n+1} – lap_n$ COM shifts as a function of post-onset laps (based on the 2649 PFs trajectories in Fig. 5A). Automatic uniform binning algorithm. **B.** The proportion of absolute shifts larger than 3 cm (edge of the center bins in A) decreases after onset but stabilizes to a non-zero value. **C-F.** Using larger shift thresholds supports the same conclusion that sizeable shifts are more frequent right after PF onset but their probability relaxes to a positive constant.
